# Gene Specific Pathogenicity Predictor for Chromatin-Remodeling BAF Complex-Associated Neurodevelopmental Disorders

**DOI:** 10.1101/2025.09.11.675179

**Authors:** Joshua Hack, Mohammad Nazim

## Abstract

Advancements in whole genome sequencing have increased the number of variants of uncertain significance (VUS) identified in patient genomes. This has created a diagnostic bottleneck for genetic counselors tasked with sifting through these variants and determining those most likely to be causative for a patient’s clinical presentation. Machine learning (ML) tools can aid in identifying pathogenic variants from VUS, but there is a need for gene-specific algorithms that predict pathogenic variants with high accuracy. To address this need, we present a workflow for developing gene-specific, ensemble-learning ML tools, that leverage outputs from other algorithms, locations of variants within the gene, and evolutionary conservation data to make a prediction of pathogenicity. Variants in *SMARCA2* and *SMARCA4* that are associated with rare neurodevelopmental diseases were used to screen 15 ML algorithms. A random forest learner was tuned to yield a final accuracy of 0.93 on holdout data. Generalizing this predictor to other BAF complex proteins resulted in a sharp decline in performance. We trained a final predictor for all genes in the study to create a predictor that identifies pathogenic variants in these BAF subunits with an accuracy of 0.91 on holdout data. This predictor specific to BAF complex proteins performs with higher accuracy and AUROC than any other predictor. The decline in performance when generalized to other proteins emphasizes the need for the gene-specific calibration of predictors. Our workflow for the development of such models provides a quick, computationally inexpensive route for improving the ML tools available to genetic counselors.

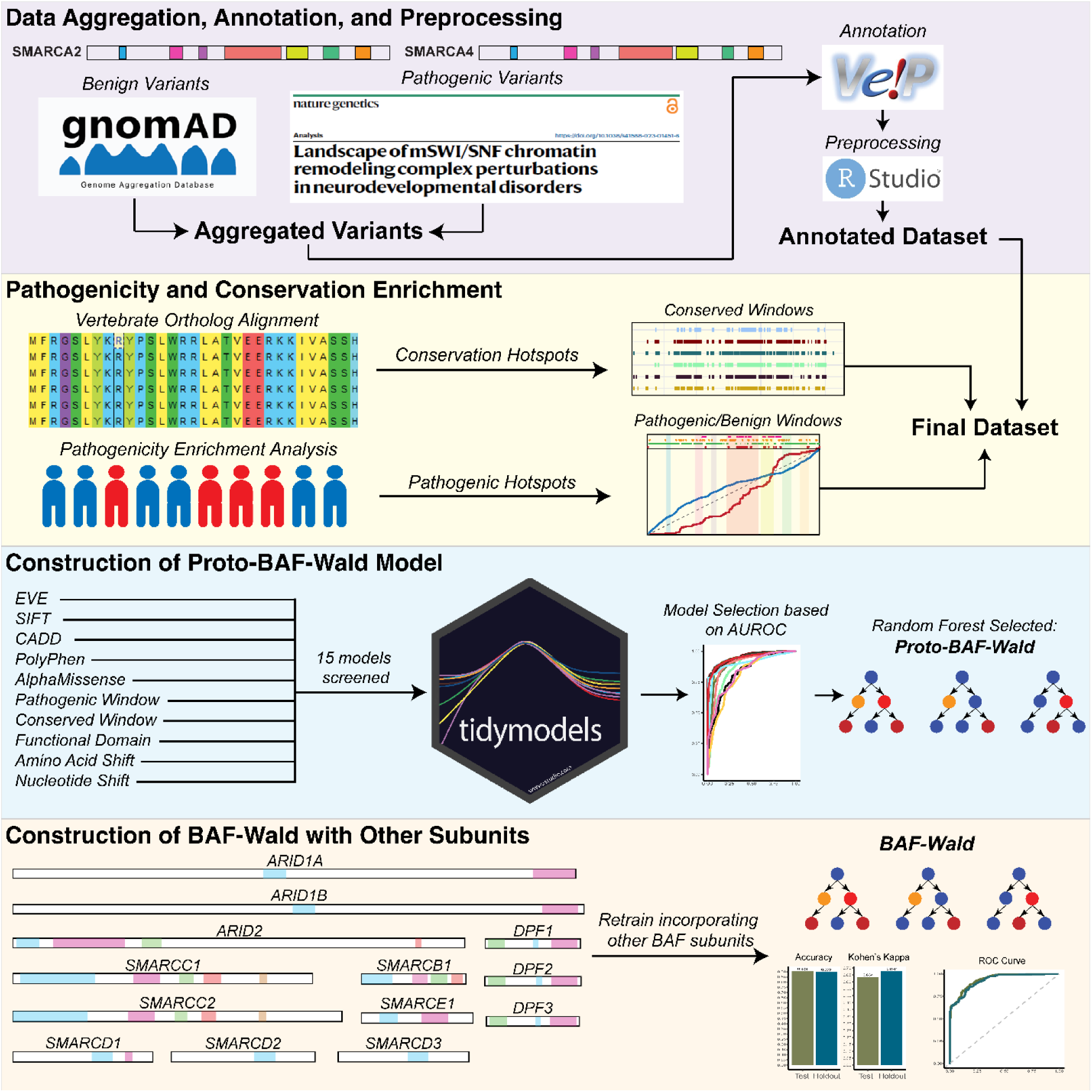
Graphical Abstract.

## Introduction

The introduction of next generation sequencing as a tool in the clinical diagnostic arsenal has resulted in rapid discovery of rare variants associated with disease. Reduction in the cost of whole exome or genome sequencing has made it possible to sequence more patients and identify variants that may be causal for their symptoms. However, these sequencing techniques often return dozens of variants of uncertain significance (VUS)^1–3^. An alternative approach involves sequencing of gene panels associated with a largely heterogeneous disease, such as epilepsy^4^ or neurodevelopmental disorders^5^. This approach reduces the number of variants to be assessed but excludes possibly pathogenic variants that are not already incorporated into those gene panels. The difficulty for genetic counselors and clinicians is collating information from different databases and literature to determine which potential VUS is the most likely causal variant, resulting in a diagnostic bottleneck^6,7^.

Over the last 20 years, several machine learning (ML) tools have been developed with the goal of predicting whether a variant is pathogenic or benign. These tools are differentiable based on which type of genomic data they are trained on. Tools such as Polyphen and CADD prioritize sequence data and how changes in sequence may translate to protein function^8,9^. Others such as SIFT and EVE are based on evolutionary conservation of nucleotide sequence^10,11^. Finally, there are some algorithms such as AlphaMissense that consider more directly how protein folding and interactions may be impacted by a variant^12^. Each of these tools have different strengths and limitations, and while they generally perform well on common variants, they struggle to accurately classify rare variants in genes that are poorly characterized^13,14^. To address this issue, Rare Exome Variant Ensemble Learner (REVEL) was developed to act as an ensemble learner that combines predictions from a variety of these tools to deliver a final verdict^15^.

Despite these advances and the variety of pathogenicity predictors available, these tools are difficult to generalize to novel variants. Thus, there is a need for gene-specific algorithms, particularly for diseases that are diverse in underlying genetic etiology. One such disease family is that of neurodevelopmental disorders (NDD), whose causes span a wide range of protein families, including voltage-gated channels, scaffold protein binding proteins, and transcription coregulators. The latter of these comprises the largest class of genes reported in DECIPHER for developmental disorder associated genes. The largest subclass contains genes encoding subunits of the BAF complex (BRG1/BRM-associated factor complex), also known as the mammalian SWI/SNF chromatin remodeling complex^16^. Developing a pathogenicity predictor for genes within this complex would greatly increase our capacity to properly identify causative variants in a substantial number of patients with NDDs.

The BAF complex harnesses ATP hydrolysis to reshape chromatin structure and regulate transcription factor accessibility to target genomic regions^17–20^. In mammals, BAF complexes are built from at least 15 subunits encoded by about 29 different genes^17^. By interchanging these subunits, cells create a variety of BAF assemblies during development, each with unique functional specificities^21–23^. We have recently shown that during neuronal development, the BAF complex reorganizes not only through subunit switching but also by incorporating alternative splice variants of one of its subunits^24,25^. A recent comprehensive analysis of sequence variants from multiple NDD datasets showed that BAF subunits harbor an unusually high frequency of mutations in these disorders^16^. Developing a pathogenicity predictor for genes encoding the BAF subunits would greatly increase our capacity to properly identify causative variants in a substantial number of patients with NDDs.

Here we present an ensemble method for developing pathogenicity prediction tools that include gene-specific features and based on clinical and evolutionary data. The algorithm (BAF-Wald) is initially trained on variants in the BAF complex subunits SMARCA2 and SMARCA4, which are mutually exclusive and necessary ATPase subunits of the complex. Following construction of the predictor, we generalize the BAF-Wald algorithm to other components of the BAF complex including other SMARC subunits and the DPF and ARID protein families. The BAF-Wald algorithm is compared against each of the predictors utilized as input features and compared against REVEL – another ensemble algorithm – and is found to outperform each of these in predicting the consequence of variants in BAF complex subunits.

## Methods

### Data collection

BAF-complex variants associated with NDD were collected from SFARI, DECIPHER, ClinVar, SPARK, SSC-ASC, DDD, LOVD, and reported sequence variants from the literature in previously published work^16^ and filtered to select genes in the SMARC subunits and DPF and ARID families. Variants were collected from gnomAD that were not present in the pathogenic database and were not annotated as pathogenic, likely pathogenic, variants of uncertain significance (VUS), or possessing conflicting pathogenicity reports to serve as benign variants^26^. All variants were split into two datasets: *Initialization,* which included *SMARCA2* and *SMARCA4,* and *Generalization,* which included the remaining genes (**Table S1**). Every variant was passed through the Variant Effect Predictor (VEP) tool hosted by Ensembl^27^ to curate predictions from previously constructed tools, namely: AlphaMissense^12^, Combine Annotation Dependent Depletion (CADD)^9^, Polymorphism Phenotyping version 2 (PolyPhen)^8^, Evolutionary Model of Variant Effect (EVE)^11^, Sorting Intolerant from Tolerant (SIFT)^10^, ClinPred^28^, and SpliceAI^29^. Performance metrics were calculated for each predictor (**Table S2**). In addition to all predictions, the variant codon position, amino acid shift, nucleotide shift, and if the variant occurred in a functional region of the protein were included as features.

### Pathogenic region identification

Regions enriched for pathogenic or benign variants-deemed to be “hot spots”-were identified as described in previous work^30^. In short, cumulative distribution lines were plotted for the variants in the patient dataset and gnomAD based on the nucleotide position within the mRNA transcript. Regions were identified to be either “flat” or “steep” depending on if there was an excess of variants across a contiguous sequence. Regions were then called lethal if both cumulative distributions were flat, complex if both were steep, pathogenic if the patient distribution was steep while the public distribution remained flat, or benign if the public distribution was steep while patient distribution was flat. Sections are termed lethal under the assumption that the absence of variants reported in either database is due to these variants in these regions being poorly tolerated. Variants included in the study were annotated as residing within a “pathogenic”, “benign”, or “other” region.

### Conservation analysis

Evolutionary conservation of each gene and biochemical properties of amino acids across vertebrates was calculated. In brief, ortholog protein sequences were downloaded from NCBI’s ortholog database and aligned in UNIPRO Gene using Kalign^31^ with a gap open penalty of 2.54 and gap extension penalty of 0.13. Alignments were loaded in MEGA 11 and sequences that were identified as low quality or predicted proteins were deleted before conducting a phylogenetic analysis. Each amino acid site was evaluated for conservation rate across collected vertebrate sequences and amino acids were classified into groups based on biochemical properties (i.e., polarity, acidity, structure, size, origin) for evaluation of these properties. Highly conserved regions (>98%) were identified using a sliding window of 10 amino acids and variants that reside in these regions were annotated as an additional feature in the study dataset.

### Modeling

To determine the ideal preprocessing and machine learning algorithm for this dataset, 15 workflows were defined using the “tidymodels” package in RStudio v4.2.2. Data from the *Initialization* subset using only *SMARCA2* and *SMARCA4* variants were subset into 60% training, 20% testing and 20% holdout validation sets, and training was done across 5 folds. Data was preprocessed generally to encode three categorical features to numerical features, remove highly correlated features, impute missing values using k-nearest neighbors (n=10), and finally normalized. Additional preprocessing recipes were constructed to include SMOTE sampling and conduct additional kNN (n=5) imputation. Algorithms that were screened and their preprocessing recipes are described in **Table S3**. All workflows were evaluated using accuracy, specificity, sensitivity and area under the receiver operating characteristic curve (AUROC).

Following screening, the random forest algorithm with no SMOTE sampling was determined to be the highest performing workflow. This workflow was tuned to determine optimal number of trees between 500 and 2000, number of variables between 2 and 5 variables, and minimum nodes between 2 and 10. The optimal parameters were selected based on AUROC and the final tuned model was fit on the training data. The tuned and fitted random forest model was evaluated on the test and holdout validation sets for accuracy and AUROC to determine if overfitting had occurred.

The tuned model was then tested on the variants in the *Generalization* dataset to determine if training on a limited subset of variants could generalize to other genes encoding subunits of the BAF complex. Performance was evaluated using accuracy and AUROC for variants subset into SMARC genes, DPF genes, and ARID genes. Following this, a new model was trained, tuned, and fit using the combined data from both the *Initialization* and *Generalization* datasets for a final tuned model able to predict the pathogenicity of variants in these three protein families.

## Results

### SMARCA2/4 have regional hotspots for pathogenicity

Surveying variants reported in the patient population and in healthy populations identified regions within *SMARCA2* and *SMARCA4* that are enriched for pathogenic or benign variants (**Figure 1**). In both *SMARCA2* and *SMARCA4*, neither pathogenic nor benign variants were randomly distributed across the protein (Anderson Darling test: SMARCA2: benign p= 8.434×10^−48^, pathogenic p= 2.52×10^−127^; SMARCA4: benign p=1.03×10^−51^, pathogenic p= 3.77×10^−30^). Notably, the regions of both genes that are enriched for pathogenic and lethal variants are the helicase domains which are responsible for ATP-dependent unwinding of DNA. Both genes have regions determined to be “complex”, regions that are enriched with both benign and pathogenic variants. This may be due to an excess of reported benign variants in the database, skewing these regions to complex when they may be more pathogenic, or could suggest these regions are tolerant to amino acid shifts that conserve biochemical properties while shifts that result in substantial biochemical changes (i.e., polarity, acidity) are more likely to be deleterious.

**Figure 1.**
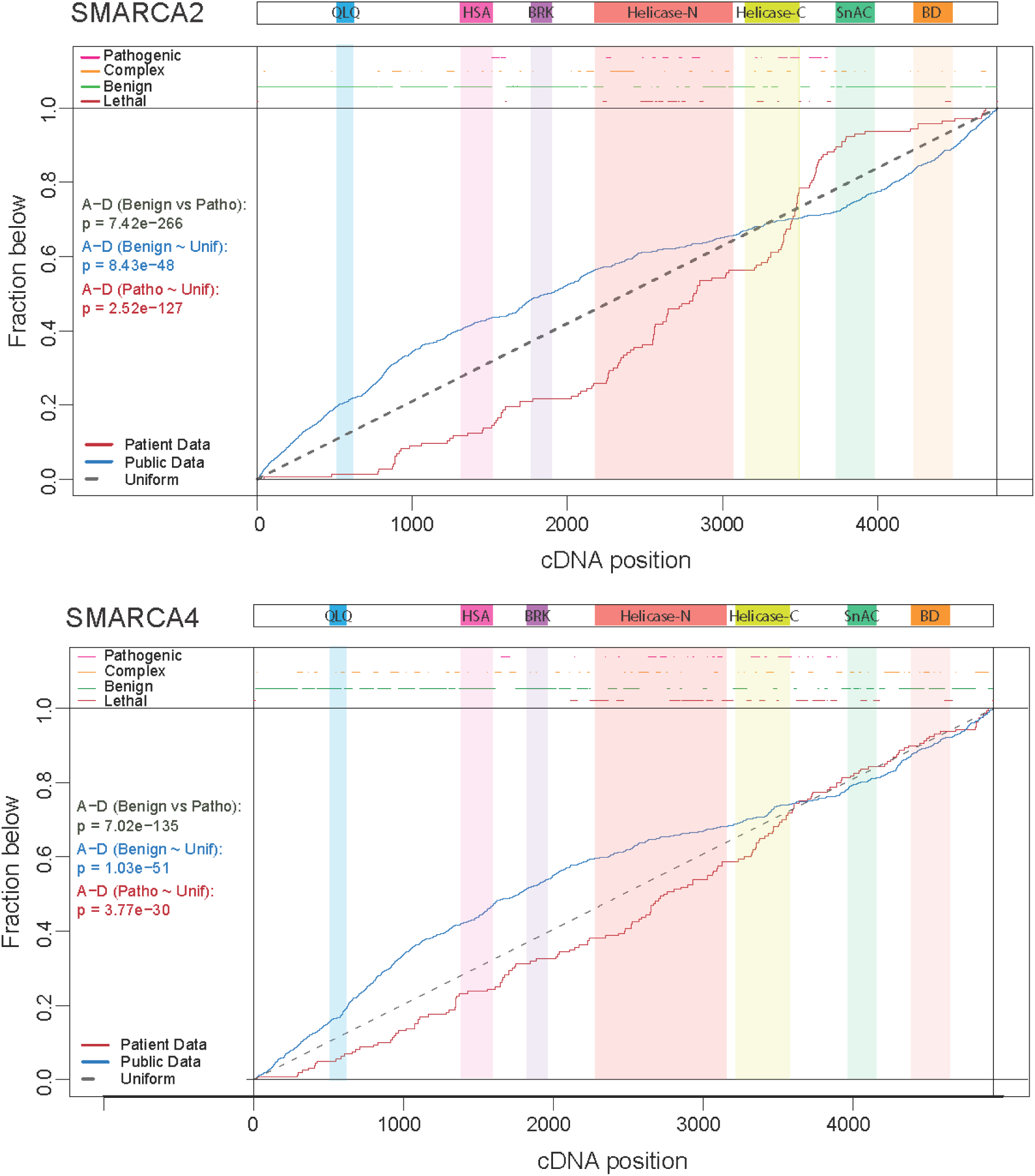
Pathogenicity plots for SMARCA2 and SMARCA4. Cumulative distribution plots of variants that are reported as benign in gnomAD (blue) and pathogenic as reported by Valencia et al. (red) for SMARCA2 and SMARCA4. Regions enriched for pathogenic or benign regions are annotated and regions depleted of any variant are annotated as “lethal”. Results from Anderson-Darling tests between the benign and pathogenic distributions and between both distributions and a uniform distribution. QLQ, Glutamine–Leucine– Glutamine motif containing domain; HSA, Helicase/SANT-associated domain; BRK, Brahma and Kismet domains; SnAC, Snf2 ATP-coupling domain; BR, Bromodomain.

### SMARCA2/4 conservation across vertebrates

As expected, both *SMARCA2* and *SMARCA4* are highly conserved across vertebrates (**Figure 2A-F**). For *SMARCA2* and *SMARCA4,* respectively, 70% and 75% of amino acids are conserved in at least 98% of vertebrates (**Figure 2C-D**). Functional domains of both proteins are highly conserved in both biochemical properties and general amino acid sequence (**Figure 2E-F**). The high conservation throughout these proteins is expected, as *SMARCA2* or *SMARCA4* are necessary components of the SWI/SNF chromatin remodeling complex. Furthermore, these two proteins are the only subunits of this complex with an ATPase domain to utilize ATP for chromatin remodeling activities^32–34^.

**Figure 2.**
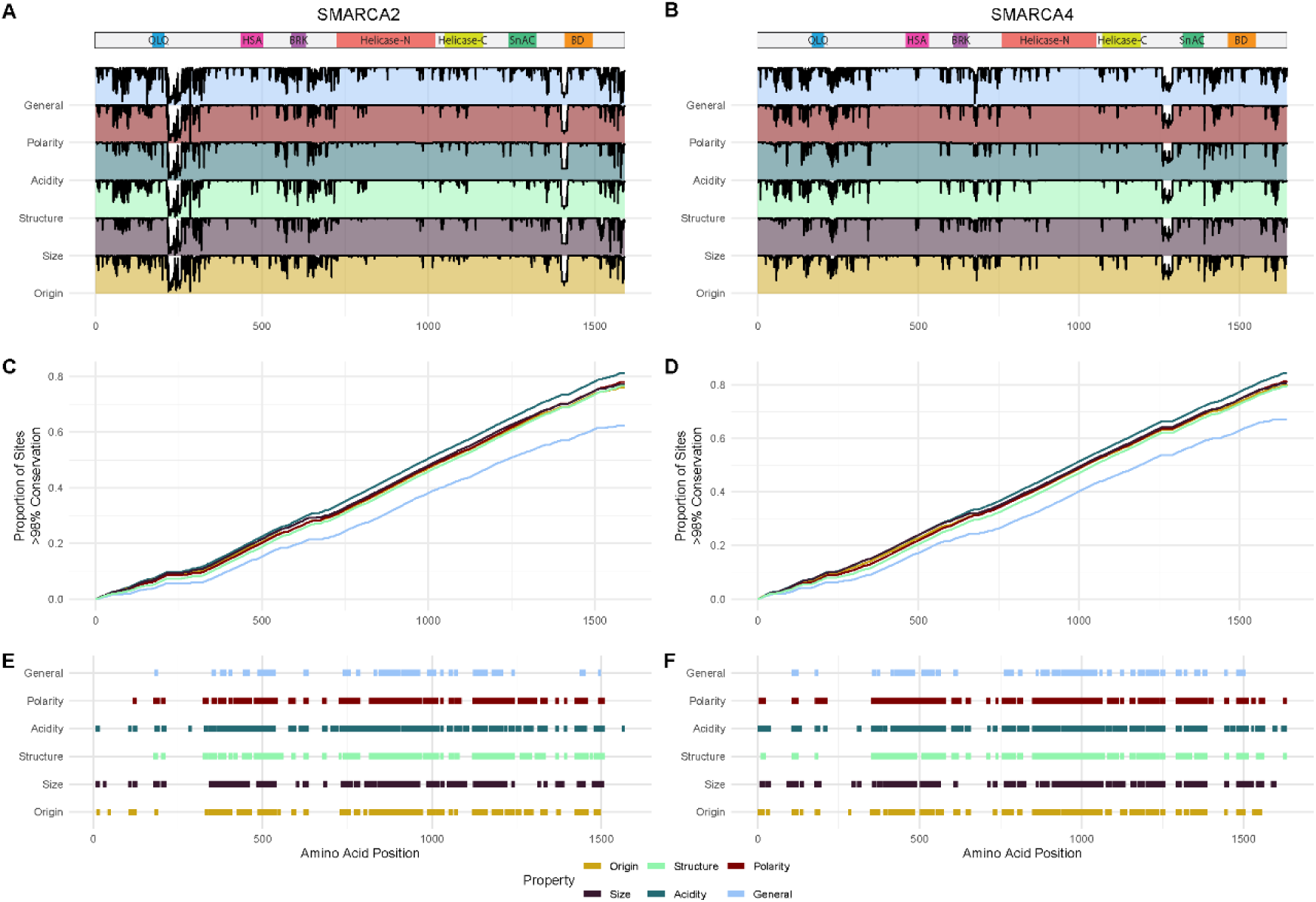
Conservation of amino acids and biochemical properties among vertebrates for SMARCA2 and SMARCA4. A-B) Ridge plots of conservation of the amino acid sequence (general), four biochemical properties (polarity, acidity, structure, size), and if the amino acid is essential or nonessential (origin). C-D) Cumulative distribution plots for protein sites with >98% conservation across vertebrates. E-F) Plot made using a sliding 10-site window to determine regions that are conserved with >98%. Bars indicate regions in which the majority of sites are conserved within the 10-site window.

### Random Forest identified as optimal algorithm

Fifteen different learner strategies were tested to identify which algorithm would perform with highest accuracy, sensitivity, specificity, and AUROC. The highest performing algorithms each rely on decision trees for predictions that vary in how the final decision is made. Random forests, which are comprised of trees that use a variable number of features within the dataset, were found to be the optimal model for maximizing all metrics (**Figure 3A-C, Table S4**). Bagged decision trees, which use the same number of features for each decision tree, also performed with high metrics, although were less accurate on average compared to random forests without SMOTE resampling (**Figure 3A**). Of the algorithms that do not utilize decision trees, a weighted logistic regression model without resampling was the highest performing. Following this, Bayesian models and neural networks were the highest performing algorithms. Support vector machines were much more variable in performance and generally performed worst out of all models, although a linear support vector machine performed with comparable accuracy to other learners (**Figure 3A-B**).

**Figure 3.**
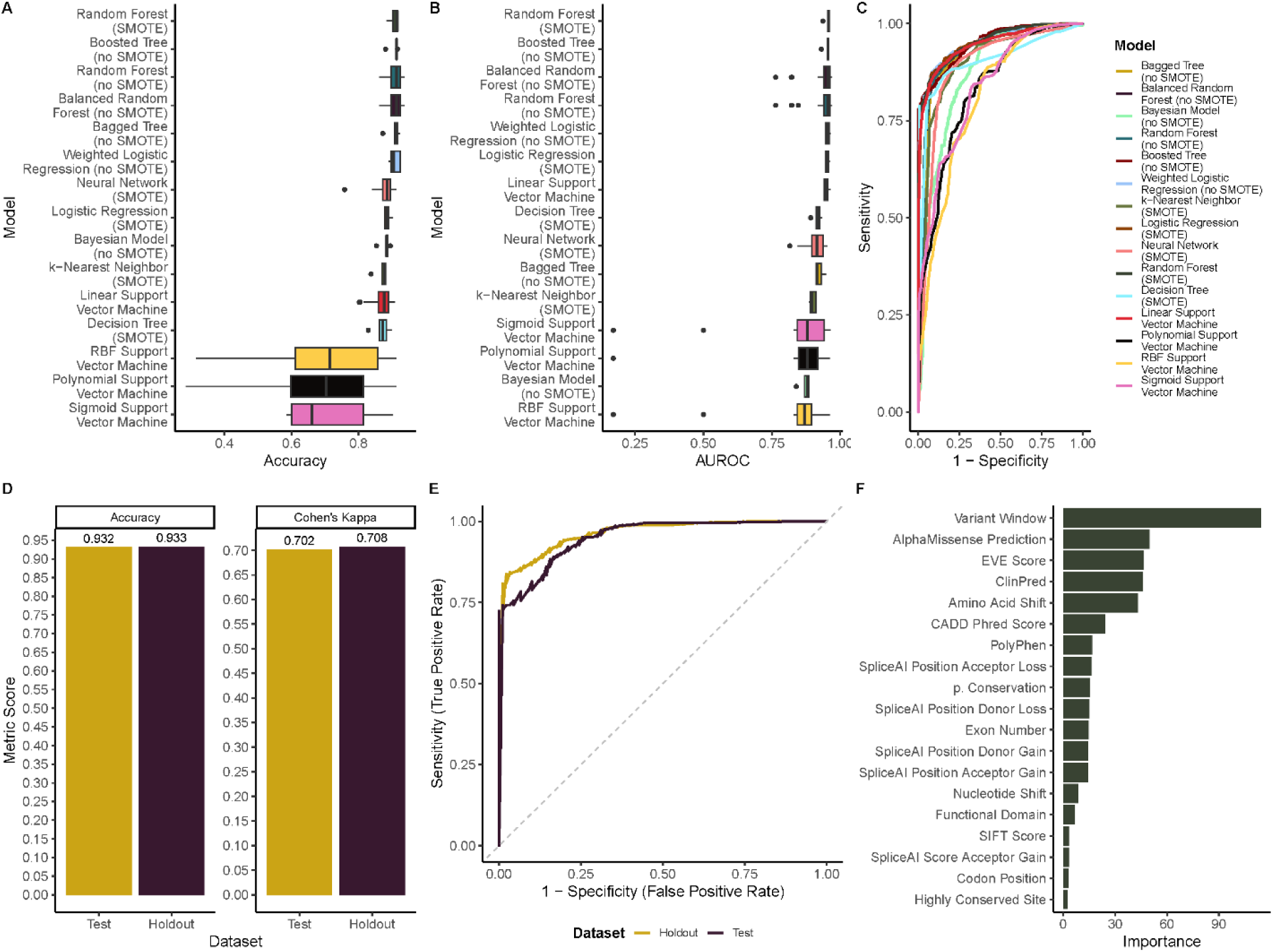
Results from machine learning modeling of *SMARCA2* and *SMARCA4*. Accuracy (A), area under the receiver operating characteristic curve (B), and receiver operating characteristic curve (C) of each of the 15 recipes screened across five folds. D) Accuracy (left) and adjusted accuracy (right) of the final tuned random forest model applied to test and holdout datasets. E) Receiver operating characteristic curves for the final model applied to test and holdout datasets. F) Feature importance for the random forest in determining whether a variant is pathogenic or benign.

Following screening, a random forest model without resampling or balancing was selected to tune and validate on holdout data. The final hyperparameters for the random forest were 1143 trees, a minimum of 3 nodes, and 4 variables (**Table S5**). Following tuning, the algorithm predicts pathogenicity with 93.2% accuracy on the test data and 93.3% accuracy on holdout validation data (**Figure 3D**, **Table S6**). Evaluation of each feature for importance in determining prediction identified the variant residing in a pathogenic variant window (defined by cumulative distribution, **Figure 1**) as the most important feature, followed by the predictions from AlphaMissense, EVE, and ClinPred, and the amino acid shift (**Figure 3F**). The tuned random forest model compares favorably with each of the prediction tools used in this study and outperforms the REVEL ensemble learner^15^ with cutoffs at both 0.5 and 0.75 as an alternative ensemble ML method (**Table S2**).

### Trained predictor does not generalize to other BAF Complex proteins

To test whether a pathogenicity predictor trained on two genes could generalize to genes encoding the other core subunits of the BAF complex, pathogenic and benign variants were collected and annotated for *SMARCB1, SMARCC1, SMARCC2, SMARCD1, SMARCD2, SMARCD3,* and *SMARCE1*. Pathogenic and benign windows were identified using the same method for *SMARCA2* and *SMARCA4* and each gene amino acid sequence was evaluated for conservation of each site across vertebrates. The plots generated from this for all proteins are presented in **Figures S1-13**. Following variant annotation using VEP, the previously tuned random forest algorithm was tested using these proteins. The performance of the algorithm substantially decreases when generalized to proteins within the same complex, with an accuracy of 0.58 and an AUROC of 0.54 (**Figure 4B-C, Table S7**). The low accuracy in this case may be due to the dearth of pathogenic variants identified in SMARC subunits outside of *SMARCA2* and *SMARCA4*, resulting in difficulty establishing pathogenic windows.

**Figure 4.**
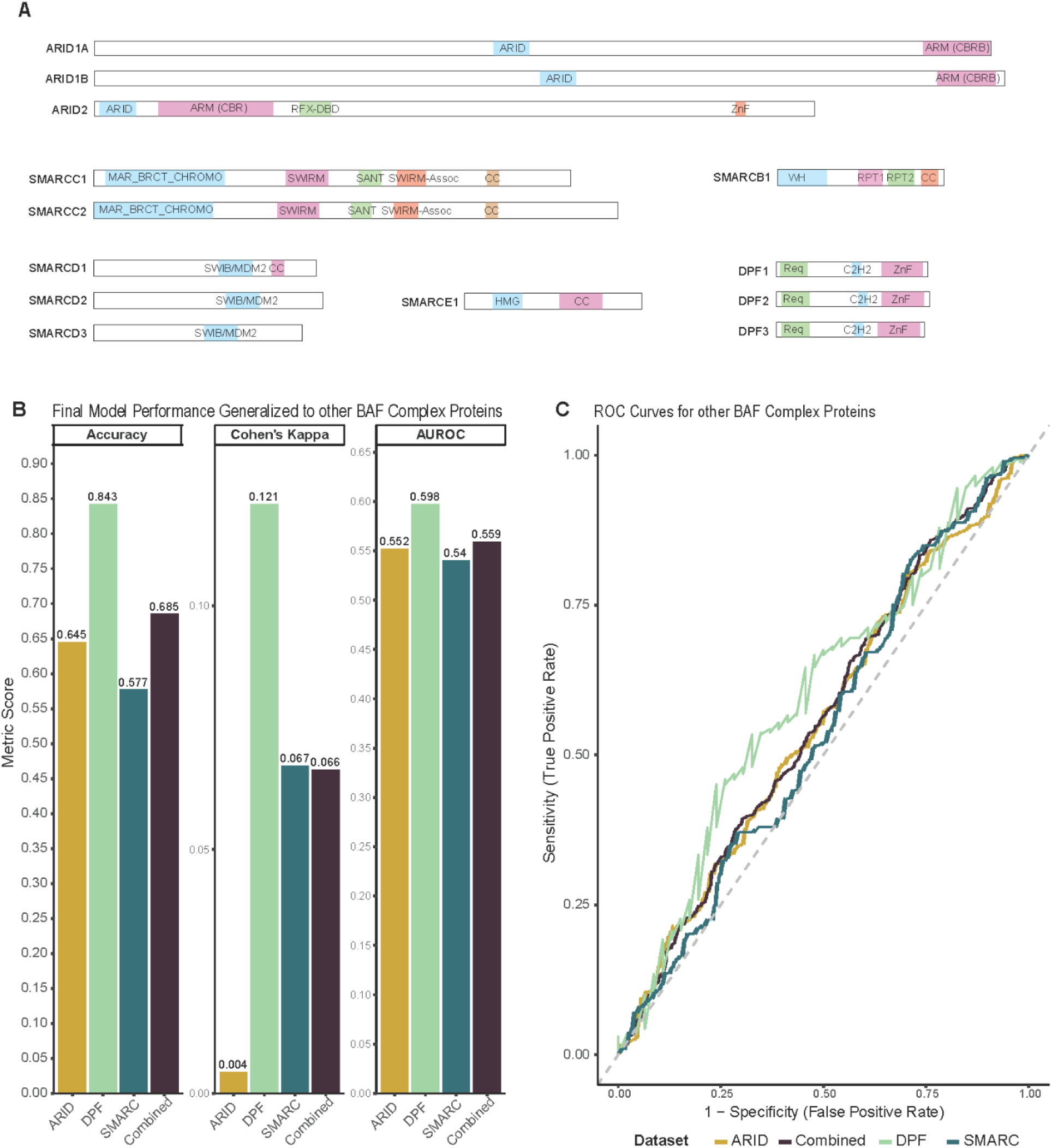
Performance metrics of the random forest model applied to other BAF complex subunits. A) Annotated protein structures for SMARC, DPF, and ARID subunits. Domain annotations and sequence locations were identified using UniProt and PAMK. B) Accuracy (left), adjusted accuracy (middle), and area under the receiver operating characteristic curve (right) for the model applied to ARID, DPF, and SMARC subunits. C) Receiver operating characteristic curves for the model applied to ARID, DPF, and SMARC subunits.

Similarly, the predictor’s generalizability to other BAF complex subunits was evaluated using variants collected for *DPF1-3, ARID1A, ARID1B,* and *ARID2.* This was done to determine if the features used in modeling were sufficient for application to genes in dissimilar structural families without the need to include variants from those genes in the training. Interestingly, the predictor performs better in both accuracy and AUROC in both the DPF and ARID families relative to the SMARC subunits. The decrease in accuracy is marginal when generalized to DPF (accuracy=0.843) while generalization to the ARID proteins results in a greater decrease in accuracy (accuracy=0.645) (**Figure 4B-C**). Despite the prominent differences in accuracy between these protein families, each subunit yields similar AUROC values (SMARC=0.54; DPF=0.598; ARID=0.552) (**Table S7**). While the results from generalizing to DPF are encouraging, normalizing the accuracy to what is expected by chance alone indicates the generalization is not as effective. In conclusion, the tuned algorithm does not generalize well to other genes within the same protein complex.

However, when compared to each of the individual predictors and REVEL, the algorithm still outperforms 7/8 models in accuracy and AUROC, although the sensitivity of the algorithm is lower than all predictors except REVEL with a pathogenicity cutoff of 0.75 (**Table S2**).

### Inclusive reconstruction of algorithm yields high performing pathogenicity predictor

Since generalization of the algorithm was ineffective in predicting pathogenic variants in other genes, the algorithm was reconstructed using all genes included in this study. The same workflow for model screening and tuning was conducted with similar results. A random forest with no SMOTE resampling was identified as the optimal algorithm, and tuning identified the ideal parameters for optimizing performance (**Figure 5A-C, Table S8**). Once tuned, the algorithm predicted pathogenicity with high accuracy in the training (0.998) (**Table S9**), test (0.903), and holdout sets (0.899), outcompeting all other predictors for the included genes (**Figure 5D-E, Table S2**). The relative importance of each feature was consistent with the algorithm trained on only *SMARCA2* and *SMARCA4* (**Figure 5F**). To ensure the model was not overly reliant on the variant residing in a pathogenic region, a final test was performed with the removal of this feature. There was no difference in performance following the removal of variant window (**Table S10-S11**). These results emphasize the need for gene-specific data and training for predictors rather than relying on generalized predictors trained on a smaller subset of genes.

**Figure 5.**
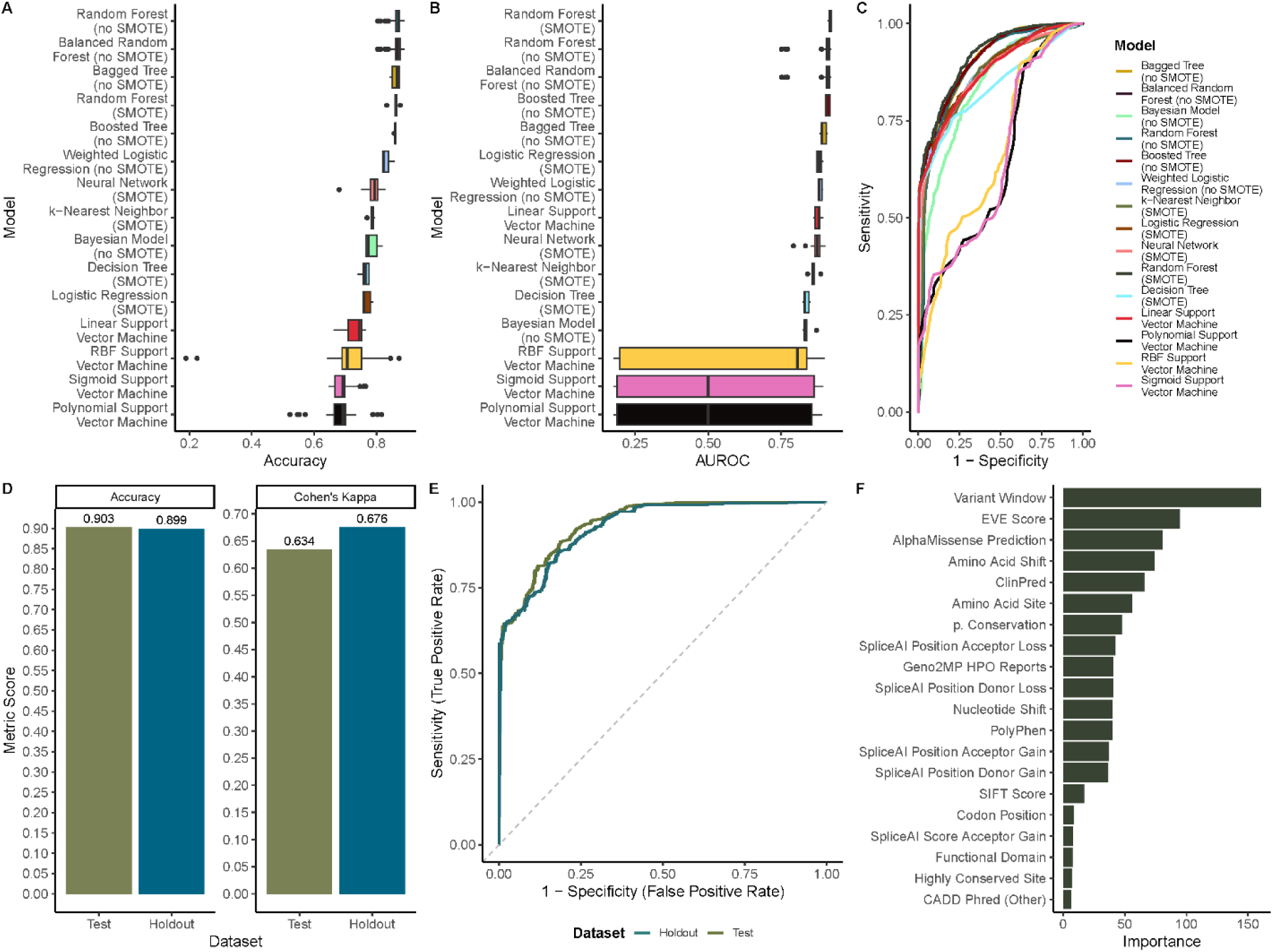
Results from machine learning modeling including all SMARC, DPF, and ARID subunits. Accuracy (A), area under the receiver operating characteristic curve (B), and receiver operating characteristic curve (C) of each of the 15 recipes screened across five folds. D) Accuracy (left) and adjusted accuracy (right) of the final tuned random forest model applied to test and holdout datasets. E) Receiver operating characteristic curves for the final model applied to test and holdout datasets. F) Feature importance for the random forest in determining whether a variant is pathogenic or benign.

## Discussion

One of the greatest challenges genetic counselors face is the interpretation of rare VUS. Over 40% of individuals that undergo sequencing will have at least one such variant reported, leaving genetic counselors to determine which VUS is causative^35^. While most of these variants are later reclassified as benign or likely benign according to the guidelines of the American College of Medical Genetics^36^, the counselor must decide which variant is pathogenic with a dearth of data. As more novel variants are identified and reported, the time needed to properly evaluate each variant results in a diagnostic bottleneck, delaying treatment that is well informed. This can have detrimental effects on the patient, particularly if the disease phenotype has a standard protocol that is contraindicated for specific diseases. One such case is SCN8A-Related Epilepsies, in which patients are detrimentally affected by treatment with the anti-seizure medication levetiracetam, the standard first-line therapeutic in pediatric epilepsies^37–39^.

A substantial amount of sequencing data indicates that mutations in genes involved in chromatin remodeling are linked to a broad spectrum of human diseases, including multiple cancers and neurodevelopmental disorders (NDDs) ^16,17,19,40–43^. A recent large-scale analysis of sequence variants across diverse NDD datasets revealed that chromatin remodeling factors carry a disproportionately high mutational burden in NDDs ^16^. Gene ontology analysis of molecular functions further showed that the most significantly disrupted processes caused by these mutations are related to transcription and chromatin regulation ^16^. Notably, the mammalian ATP-dependent chromatin remodeling SWI/SNF complex, also known as the BRG1-Associated Factor (BAF) complex, harbors the greatest number of NDD-associated *de novo* missense and protein-truncating variants (PTVs), most of which are classified as pathogenic or likely pathogenic ^16^. Many of these mutations map to highly conserved regions and functional domains of BAF subunits ^16^. However, the molecular mechanisms of pathogenicity are poorly understood, largely due to the absence of gene-specific pathogenicity prediction models. Elucidating how genetic mutations disrupt chromatin remodeling complexes is therefore critical for understanding the molecular basis of neurodevelopmental disorders and for advancing the development of targeted therapeutic strategies.

There are tools available that aid in the classification of variants as pathogenic or benign, but many of these are constructed with common variants, limiting their utility for rare disease applications ^7,13^. Other tools are biased toward genes that are well characterized in vertebrates or have an excess of functional or structural studies ^44^. These limitations lead to lack of generalizability and poor performance for novel variants in genes that are more recently characterized or less frequently studied. One approach to overcoming these limitations is to use ensemble machine learning, constructing a classifier that uses the outputs of other tools to come to a final decision. However, even these ensembled learners are poorly generalized to rare variants. In light of this, some have remarked on the need for gene-specific predictors. These gene-specific predictors present their own challenges, such as limited clinical data on many genes, poor annotations on variants without functional studies or case studies that link variants to a disease, and the vast number of genes that would require specific predictors.

In an attempt to streamline the process of generating these gene-specific classifiers, we leveraged ensemble learning with gene-specific analysis for regions enriched with pathogenic and benign variants, reports of phenotype associated with variants of uncertain significance in the Geno2MP dataset, and analysis of conserved variants and regions. This random forest model (*BAF-Wald*) was initially trained on two SMARC proteins before being generalized to other proteins within the BAF complex with the goal of predicting pathogenic variants that cause neurodevelopmental disorders. The remainder of this discussion will evaluate the model’s performance compared to other tools, and discussion of the feasibility of applying this workflow to other genes.

### BAF-Wald compared to other predictors

Previously constructed pathogenicity predictors utilized a variety of data types and sources for their training. This study uses seven different predictors as input features for ensemble learning: AlphaMissense^12^, Combine Annotation Dependent Depletion (CADD)^9^, Polymorphism Phenotyping version 2 (PolyPhen)^8^, Evolutionary Model of Variant Effect (EVE) ^11^, Sorting Intolerant from Tolerant (SIFT)^10^, ClinPred^28^, and SpliceAI^29^. The six former predictors are tasked with predicting pathogenicity from missense variants while SpliceAI is used to predict if a variant will cause gain or loss of splice donor or acceptor sites. All models are useful for clinical purposes, but each has its own limitations due to the data used to train the predictor.

AlphaMissense is the newest of these predictors, being first published in 2023, and is trained on a combination of evolutionary data and predictions from AlphaFold^12,45^ regarding the variant’s impact on protein structure. This predictor excels in proteins with extensive annotations of structural components, a history of protein structure analysis through crystallography, and high-quality sequencing of diverse vertebrates. However, proteins without this study profile have poor performance with AlphaMissense, due to low confidence predictions of protein folding or limited species to effectively evaluate conservation.

CADD utilizes over 60 genomic annotations such as conservation, epigenetics, and transcription factor binding sites. This algorithm also combines real variants with simulated variants in its training. CADD struggles with regions that are poorly annotated, exacerbated by the number of annotations necessary for CADD to perform well.

PolyPhen-2 relies on sequence data, structural features, and evolutionary conservation and was trained on datasets such as HuVarBase and HumDiv^46^. This algorithm relies strongly on variants that are already annotated as pathogenic in clinical settings, leading to novel variants being missed if PolyPhen-2 is used to filter out variants identified in clinical settings. Since this model relies heavily on two datasets with similar variant makeup, the prediction may also be confounded by the collinearity between the datasets.

EVE employs an unsupervised machine learning approach to evolutionary conservation data. This model limits any bias that might be inherent to supervised approaches and identifies patterns that may have been missed in a purely supervised approach. However, EVE is trained to detect variants that impact overall fitness, which may miss pathogenic variants that have latent development or have little impact on the organism’s reproductive fitness while still having a clinical phenotype. ClinPred is the final ML tool that predicts pathogenicity and is trained on clinical variant data with genomic annotations using variants reported in ClinVar up to 2016. This model excels with genetic diseases with established populations but is limited in generalizing to genes with novel disease links and may be confounded by variants being misidentified as causal in some diseases.

The two remaining tools used in this study (SIFT and SpliceAI) are not strictly ML predictors of pathogenicity. SIFT is a probabilistic approach first published in 2003 that uses sequence homology and conservation across species. This predictor generally has a high false positive rate and preferentially classifies functionally neutral variants as pathogenic, particularly in highly conserved genes. Additionally, all predictors that rely on evolutionary conservation are sensitive to genes in which alternative splicing or exon skipping may occur. These genes have multiple reference sequences and depending on which sequence is used it could lead to different results. SpliceAI does not explicitly predict pathogenicity of missense variants but rather predicts whether a variant could disrupt splicing dynamics. This tool performs well with its given task but is restricted to regions surrounding splice sites for predictions and cannot assess if a variant is likely to be pathogenic if it does not cause aberrant splicing. However, it is still a valuable tool in clinical settings for determining if a variant is likely to have deleterious effects.

Each of these tools on its own is able to inform some aspects of whether a variant is likely to be pathogenic or not. However, each has limitations that decrease the confidence of genetic counselors and clinicians in identifying whether a VUS is indeed the causal variant. Thus, genetic counselors are tasked with deciding which of these tools are most reliable or if the majority of models are reliable in their output. BAF-Wald attempts to streamline this process by querying all 7 of these algorithms, adding additionally evolutionary and clinical analysis, and providing a consensus decision based on the ensemble of algorithms. To determine the success of the algorithm, it must perform better than all of the included algorithms performed independently. Indeed, BAF-Wald performs with higher accuracy, specificity, and AUROC than all models in predicting the pathogenicity of variants in *SMARC, DPF,* and *ARID* proteins (**Table S2 & S9**). The next closest individual algorithm is EVE (0.707) and AUROC (0.768) in terms of accuracy, and CADD in terms of specificity (0.906). Impressively, when BAF-Wald was initially trained on *SMARCA2/4* and generalized to other proteins, it performed with the highest specificity (0.999) and second highest accuracy (0.630) and AUROC (0.762), being second to CADD (0.654) and EVE (0.768) for each, respectively. This relative performance lends further support to the hypothesis that ensemble learning using previously developed pathogenicity predictors results in greater cumulative performance than any of the individual predictors.

This hypothesis was already supported by improvements in performance over any individual predictor by REVEL, a previously published ensemble learner that leverages outputs from 18 different scores, 10 of which are functional and 8 are conservation. This algorithm performs well with variants that have allele frequencies <0.5% and performs better than individual predictors in general use for variants throughout the genome. However, previous work has shown that REVEL predictions are of limited value for disease specific classification of VUS as pathogenic or benign^13^. Similarly, here we find that REVEL performs with lower metrics at pathogenicity thresholds of both 0.75 (accuracy= 0.613; sensitivity=0.073; specificity= 0.943; AUROC estimate= 0.561) and 0.5 (accuracy= 0.587; sensitivity= 0.263; specificity= 0.787; AUROC estimate= 0.560) (**Table S2**). BAF-Wald prior to retraining using DPF, SMARC, and ARID proteins generalizes to these with greater accuracy, AUROC, and specificity than both thresholds, while maintaining higher sensitivity than pathogenicity thresholds of 0.75. This disparity in performance highlights the need for disease– and gene-specific classifiers, trained using clinical, evolutionary, and functional data.

### Generalization of workflow to other genes

One of the greatest challenges in developing gene-specific pathogenicity predictors is the time and data necessary to construct one that is accurate. This study attempts to limit this challenge by adding only two gene-specific features requiring substantial data: pathogenic regions based on clinical variants and regions of the gene that are highly conserved across all vertebrates. The former requires substantial variants that have been annotated as benign and pathogenic or reported in case or cohort studies as associated with a disease. This requirement means that predictors built in this way will not necessarily classify variants with high accuracy in diseases with small affected populations. However, this workflow appears to generalize to other proteins in the same complex better than alternative classifiers, with comparable accuracy to CADD and EVE, higher AUROC, and greater specificity, limiting the number of false positives. This allows for a lower confidence prediction of pathogenicity in early stages following description of a disease, with improvements as more pathogenic variants are reported. The concern of decreased performance without a large sample of pathogenic variants is also addressed by removing the variant window feature in this study, resulting in a marginal decrease in performance.

Overall, this workflow for construction of gene-specific pathogenicity predictors succeeds in quickly developing an effective predictor with limited clinical data. These predictors are more readily generalized to other proteins within the same complex of the original predictor and genes that operate in similar pathways or complexes than previously described predictors. This study highlights the utility of ensemble-based machine learning for leveraging the strengths of each model to account for limitations either in data source or methodology in other models. Furthermore, this workflow is easily applied to new genes and the model’s performance is robust to the incorporation of many genes, since the affected gene is not considered as an input feature in the ensembled model.

This study has its limitations however, particularly in the enrichment of benign variants that may artificially increase the accuracy of the model. This concern is mitigated by the success of a balanced random forest and random forest model with SMOTE preprocessing, both of which performed similarly to a random forest without SMOTE resampling. There may also be circularity between the incorporated algorithms and additional features added but given the relatively low importance of added features for decision by the random forest, it is reasonable to conclude that the underlying logic of the algorithm is driven by the ensembled models themselves. Another limitation of this algorithm is the low specificity and high prediction of false negatives. However, further work to incorporate other algorithms and metrics not included in this study may reduce the number of false negatives, and additional follow-up by genetic counselors and researchers further limit the risk of false negatives. Finally, this algorithm requires further testing, both for generalizing to other proteins and to validate its performance on novel data not used in this study. As such, this study serves to outline an effective framework for developing gene-specific pathogenicity predictors, but the BAF-Wald model itself requires further tuning before being utilized in clinical or academic capacities.

A large contributor to the success of BAF-Wald is the combined decision-making of the ensemble of algorithms it is trained on. Additionally, BAF-Wald leverages clinical variant data, evolutionary conservation of sequence, and biochemical features of the protein itself. This synthesizes the most important features in determining if a variant is likely to disrupt protein structure. While BAF-Wald itself does not generalize to genes not yet incorporated into the algorithm, once a new gene is incorporated the high performance of the algorithm returns. This provides a source of optimism for continuing expansion of BAF-Wald to incorporate other genes in the BAF complex, eventually moving toward all genes associated with rare NDDs. Eventually, this algorithm may develop a vast ensemble of gene-specific training and become better suited for clinical use for most rare diseases.

## Supplemental Information

Supplemental information includes 13 figures and 11 tables.

## Acknowledgements

This work was supported by National Institutes of Health grant 1R21NS140948 to M.N. We thank Dr. Douglas Black for assistance in editing the manuscript for clarity.

## Data availability statement

All relevant data and code for replication are available at 10.5281/zenodo.16929854. The pathogenic variants used in this study were previously published under a Creative Commons Attribution 4.0 International License in Valencia et al. 2023 (doi: 10.1038/s41588-023-01451-6). All benign variants were collected from gnomAD.

## Figures

**Supplementary Figures 1-13. Pathogenicity and conservation plots for each BAF subunit included in the study.** A) Cumulative distribution plots of variants that are reported as benign in gnomAD (blue) and pathogenic as reported by Valencia et al. (red). Regions enriched for pathogenic or benign regions are annotated and regions depleted of any variant are annotated as “lethal”. Results from Anderson-Darling tests between the benign and pathogenic distributions and between both distributions and a uniform distribution. B) Ridge plots of conservation of the amino acid sequence (general), four biochemical properties (polarity, acidity, structure, size), and if the amino acid is essential or nonessential (origin). C) Cumulative distribution plots for protein sites with >98% conservation across vertebrates. D) Plot made using a sliding 10-site window to determine regions that are conserved with >98%. Bars indicate regions in which the majority of sites are conserved within the 10-site window.

## References

1. Federici, G., and Soddu, S. (2020). Variants of uncertain significance in the era of high-throughput genome sequencing: a lesson from breast and ovary cancers. J. Exp. Clin. Cancer Res. 39, 46. 10.1186/s13046-020-01554-6.

2. Lincoln, S.E., Hambuch, T., Zook, J.M., Bristow, S.L., Hatchell, K., Truty, R., Kennemer, M., Shirts, B.H., Fellowes, A., Chowdhury, S., et al. (2021). One in seven pathogenic variants can be challenging to detect by NGS: an analysis of 450,000 patients with implications for clinical sensitivity and genetic test implementation. Genet. Med. 23, 1673–1680. 10.1038/s41436-021-01187-w.

3. Stoller, J.K. (2018). The Challenge of Rare Diseases. Chest 153, 1309–1314. 10.1016/j.chest.2017.12.018.

4. Fazenbaker, A.C., Munro, C.D., Carlson, J.C., Durst, A.L., and Vento, J.M. (2024). Epilepsy panel testing criteria: A clinical assessment. J. Genet. Couns. 33, 352–360. 10.1002/jgc4.1732.

5. Mellone, S., Puricelli, C., Vurchio, D., Ronzani, S., Favini, S., Maruzzi, A., Peruzzi, C., Papa, A., Spano, A., Sirchia, F., et al. (2022). The Usefulness of a Targeted Next Generation Sequencing Gene Panel in Providing Molecular Diagnosis to Patients With a Broad Spectrum of Neurodevelopmental Disorders. Front. Genet. 13, 875182. 10.3389/fgene.2022.875182.

6. Fehr, A., and Prütz, F. (2023). Rare diseases: a challenge for medicine and public health. J. Health Monit. 8, 3–6. 10.25646/11826.

7. Burke, W., Parens, E., Chung, W.K., Berger, S.M., and Appelbaum, P.S. (2022). The challenge of genetic variants of uncertain clinical significance: A narrative review. Ann. Intern. Med. 175, 994–1000. 10.7326/M21-4109.

8. Adzhubei, I., Jordan, D.M., and Sunyaev, S.R. (2013). Predicting Functional Effect of Human Missense Mutations Using PolyPhen-2. Curr. Protoc. Hum. Genet. Editor. Board Jonathan Haines Al 0 7, Unit7.20. 10.1002/0471142905.hg0720s76.

9. Rentzsch, P., Witten, D., Cooper, G.M., Shendure, J., and Kircher, M. (2019). CADD: predicting the deleteriousness of variants throughout the human genome. Nucleic Acids Res. 47, D886–D894. 10.1093/nar/gky1016.

10. Ng, P.C., and Henikoff, S. (2003). SIFT: predicting amino acid changes that affect protein function. Nucleic Acids Res. 31, 3812–3814.

11. Frazer, J., Notin, P., Dias, M., Gomez, A., Min, J.K., Brock, K., Gal, Y., and Marks, D.S. (2021). Disease variant prediction with deep generative models of evolutionary data. Nature 599, 91–95. 10.1038/s41586-021-04043-8.

12. Cheng, J., Novati, G., Pan, J., Bycroft, C., Žemgulytė, A., Applebaum, T., Pritzel, A., Wong, L.H., Zielinski, M., Sargeant, T., et al. (2023). Accurate proteome-wide missense variant effect prediction with AlphaMissense. Science 381, eadg7492. 10.1126/science.adg7492.

13. Fiorini, M.R., Dilliott, A.A., and Farhan, S.M.K. (2023). Evaluating the Utility of REVEL and CADD for Interpreting Variants in Amyotrophic Lateral Sclerosis Genes. Hum. Mutat. 2023, 8620557. 10.1155/2023/8620557.

14. Zhang, X., Walsh, R., Whiffin, N., Buchan, R., Midwinter, W., Wilk, A., Govind, R., Li, N., Ahmad, M., Mazzarotto, F., et al. (2021). Disease-specific variant pathogenicity prediction significantly improves variant interpretation in inherited cardiac conditions. Genet. Med. 23, 69–79. 10.1038/s41436-020-00972-3.

15. Ioannidis, N.M., Rothstein, J.H., Pejaver, V., Middha, S., McDonnell, S.K., Baheti, S., Musolf, A., Li, Q., Holzinger, E., Karyadi, D., et al. (2016). REVEL: An Ensemble Method for Predicting the Pathogenicity of Rare Missense Variants. Am. J. Hum. Genet. 99, 877– 885. 10.1016/j.ajhg.2016.08.016.

16. Valencia, A.M., Sankar, A., van der Sluijs, P.J., Satterstrom, F.K., Fu, J., Talkowski, M.E., Vergano, S.A.S., Santen, G.W.E., and Kadoch, C. (2023). Landscape of mSWI/SNF chromatin remodeling complex perturbations in neurodevelopmental disorders. Nat. Genet. 55, 1400–1412. 10.1038/s41588-023-01451-6.

17. Kadoch, C., and Crabtree, G.R. (2015). Mammalian SWI/SNF chromatin remodeling complexes and cancer: Mechanistic insights gained from human genomics. Sci. Adv. 1, e1500447. 10.1126/sciadv.1500447.

18. Ronan, J.L., Wu, W., and Crabtree, G.R. (2013). From neural development to cognition: unexpected roles for chromatin. Nat. Rev. Genet. 14, 347–359. 10.1038/nrg3413.

19. Sokpor, G., Xie, Y., Rosenbusch, J., and Tuoc, T. (2017). Chromatin Remodeling BAF (SWI/SNF) Complexes in Neural Development and Disorders. Front. Mol. Neurosci. 10, 243. 10.3389/fnmol.2017.00243.

20. Narayanan, R., and Tuoc, T.C. (2014). Roles of chromatin remodeling BAF complex in neural differentiation and reprogramming. Cell Tissue Res. 356, 575–584. 10.1007/s00441-013-1791-7.

21. Lessard, J., Wu, J.I., Ranish, J.A., Wan, M., Winslow, M.M., Staahl, B.T., Wu, H., Aebersold, R., Graef, I.A., and Crabtree, G.R. (2007). An Essential Switch in Subunit Composition of a Chromatin Remodeling Complex during Neural Development. Neuron 55, 201–215. 10.1016/j.neuron.2007.06.019.

22. Staahl, B.T., Tang, J., Wu, W., Sun, A., Gitler, A.D., Yoo, A.S., and Crabtree, G.R. (2013). Kinetic Analysis of npBAF to nBAF Switching Reveals Exchange of SS18 with CREST and Integration with Neural Developmental Pathways. J. Neurosci. 33, 10348–10361. 10.1523/JNEUROSCI.1258-13.2013.

23. Braun, S.M.G., Petrova, R., Tang, J., Krokhotin, A., Miller, E.L., Tang, Y., Panagiotakos, G., and Crabtree, G.R. (2021). BAF subunit switching regulates chromatin accessibility to control cell cycle exit in the developing mammalian cortex. Genes Dev. 35, 335–353. 10.1101/gad.342345.120.

24. Nazim, M., Lin, C.-H., Feng, A.-C., Xiao, W., Yeom, K.-H., Li, M., Daly, A.E., Tan, X., Vu, H., Ernst, J., et al. (2024). Alternative splicing of a chromatin modifier alters the transcriptional regulatory programs of stem cell maintenance and neuronal differentiation. Cell Stem Cell 31, 754–771.e6. 10.1016/j.stem.2024.04.001.

25. Nazim, M. (2024). Post-transcriptional regulation of the transcriptional apparatus in neuronal development. Front. Mol. Neurosci. 17, 1483901. 10.3389/fnmol.2024.1483901.

26. Karczewski, K.J., Francioli, L.C., Tiao, G., Cummings, B.B., Alföldi, J., Wang, Q., Collins, R.L., Laricchia, K.M., Ganna, A., Birnbaum, D.P., et al. (2020). The mutational constraint spectrum quantified from variation in 141,456 humans. Nature 581, 434–443. 10.1038/s41586-020-2308-7.

27. Harrison, P.W., Amode, M.R., Austine-Orimoloye, O., Azov, A.G., Barba, M., Barnes, I., Becker, A., Bennett, R., Berry, A., Bhai, J., et al. (2024). Ensembl 2024. Nucleic Acids Res. 52, D891–D899. 10.1093/nar/gkad1049.

28. Alirezaie, N., Kernohan, K.D., Hartley, T., Majewski, J., and Hocking, T.D. (2018). ClinPred: Prediction Tool to Identify Disease-Relevant Nonsynonymous Single-Nucleotide Variants. Am. J. Hum. Genet. 103, 474–483. 10.1016/j.ajhg.2018.08.005.

29. Jaganathan, K., Kyriazopoulou Panagiotopoulou, S., McRae, J.F., Darbandi, S.F., Knowles, D., Li, Y.I., Kosmicki, J.A., Arbelaez, J., Cui, W., Schwartz, G.B., et al. (2019). Predicting Splicing from Primary Sequence with Deep Learning. Cell 176, 535–548.e24. 10.1016/j.cell.2018.12.015.

30. Encinas, A.C., Watkins, J.C., Longoria, I.A., Johnson, J.P., and Hammer, M.F. (2020). Variable patterns of mutation density among NaV1.1, NaV1.2 and NaV1.6 point to channel-specific functional differences associated with childhood epilepsy. PloS One 15, e0238121. 10.1371/journal.pone.0238121.

31. Lassmann, T., and Sonnhammer, E.L. (2005). Kalign – an accurate and fast multiple sequence alignment algorithm. BMC Bioinformatics 6, 298. 10.1186/1471-2105-6-298.

32. Mashtalir, N., D’Avino, A.R., Michel, B.C., Luo, J., Pan, J., Otto, J.E., Zullow, H.J., McKenzie, Z.M., Kubiak, R.L., St. Pierre, R., et al. (2018). Modular Organization and Assembly of SWI/SNF Family Chromatin Remodeling Complexes. Cell 175, 1272–1288.e20. 10.1016/j.cell.2018.09.032.

33. He, S., Wu, Z., Tian, Y., Yu, Z., Yu, J., Wang, X., Li, J., Liu, B., and Xu, Y. (2020). Structure of nucleosome-bound human BAF complex. Science 367, 875–881. 10.1126/science.aaz9761.

34. Mashtalir, N., Suzuki, H., Farrell, D.P., Sankar, A., Luo, J., Filipovski, M., D’Avino, A.R., St. Pierre, R., Valencia, A.M., Onikubo, T., et al. (2020). A Structural Model of the Endogenous Human BAF Complex Informs Disease Mechanisms. Cell 183, 802–817.e24. 10.1016/j.cell.2020.09.051.

35. Chen, E., Facio, F.M., Aradhya, K.W., Rojahn, S., Hatchell, K.E., Aguilar, S., Ouyang, K., Saitta, S., Hanson-Kwan, A.K., Capurro, N.N., et al. (2023). Rates and Classification of Variants of Uncertain Significance in Hereditary Disease Genetic Testing. JAMA Netw. Open 6, e2339571. 10.1001/jamanetworkopen.2023.39571.

36. Richards, S., Aziz, N., Bale, S., Bick, D., Das, S., Gastier-Foster, J., Grody, W.W., Hegde, M., Lyon, E., Spector, E., et al. (2015). Standards and guidelines for the interpretation of sequence variants: a joint consensus recommendation of the American College of Medical Genetics and Genomics and the Association for Molecular Pathology. Genet. Med. 17, 405–424. 10.1038/gim.2015.30.

37. Hack, J.B., Horning, K., Short, D.M.J., Schreiber, J.M., Watkins, J.C., and Hammer, M.F. (2023). Distinguishing Loss-of-Function and Gain-of-Function SCN8A Variants Using a Random Forest Classification Model Trained on Clinical Features. Neurol. Genet. 9. 10.1212/NXG.0000000000200060.

39. Hammer, M.F., Xia, M., and Schreiber, J.M. (2023). SCN8A-Related Epilepsy and/or Neurodevelopmental Disorders (University of Washington, Seattle).

39. Arican, P., Gencpinar, P., Cavusoglu, D., and Olgac Dundar, N. (2018). Levetiracetam monotherapy for the treatment of infants with epilepsy. Seizure 56, 73–77. 10.1016/j.seizure.2018.02.006.

40. Bögershausen, N., and Wollnik, B. (2018). Mutational Landscapes and Phenotypic Spectrum of SWI/SNF-Related Intellectual Disability Disorders. Front. Mol. Neurosci. 11, 252. 10.3389/fnmol.2018.00252.

41. Kadoch, C., Hargreaves, D.C., Hodges, C., Elias, L., Ho, L., Ranish, J., and Crabtree, G.R. (2013). Proteomic and bioinformatic analysis of mammalian SWI/SNF complexes identifies extensive roles in human malignancy. Nat. Genet. 45, 592–601. 10.1038/ng.2628.

42. St. Pierre, R., and Kadoch, C. (2017). Mammalian SWI/SNF complexes in cancer: emerging therapeutic opportunities. Curr. Opin. Genet. Dev. 42, 56–67. 10.1016/j.gde.2017.02.004.

43. Hodges, C., Kirkland, J.G., and Crabtree, G.R. (2016). The Many Roles of BAF (mSWI/SNF) and PBAF Complexes in Cancer. Cold Spring Harb. Perspect. Med. 6, a026930. 10.1101/cshperspect.a026930.

44. Gunning, A.C., Fryer, V., Fasham, J., Crosby, A.H., Ellard, S., Baple, E.L., and Wright, C.F. (2020). Assessing performance of pathogenicity predictors using clinically relevant variant datasets. J. Med. Genet. 58, 547. 10.1136/jmedgenet-2020-107003.

45. McDonald, E.F., Oliver, K.E., Schlebach, J.P., Meiler, J., and Plate, L. (2024). Benchmarking AlphaMissense pathogenicity predictions against cystic fibrosis variants. PloS One 19, e0297560. 10.1371/journal.pone.0297560.

46. Ganesan, K., Kulandaisamy, A., Binny Priya, S., and Gromiha, M.M. (2019). HuVarBase: A human variant database with comprehensive information at gene and protein levels. PloS One 14, e0210475. 10.1371/journal.pone.0210475.

