## Supplemental Figures for "Gene Specific Pathogenicity Predictor for Chromatin-Remodeling BAF Complex-Associated Neurodevelopmental Disorders"

### SMARCB1

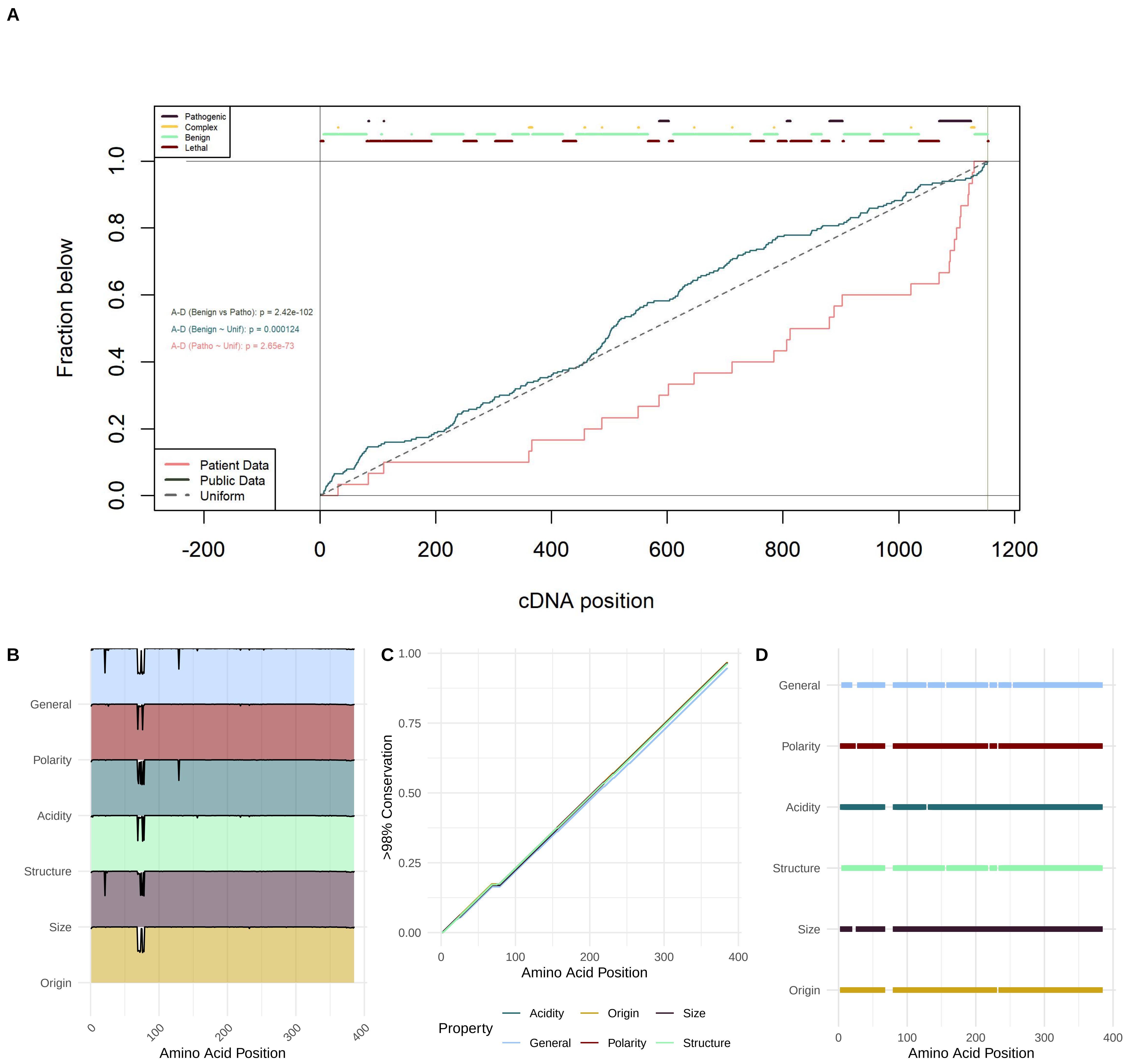
A
Pathogenic
Complex
Benign
Lethal
Fraction below
A-D (Benign vs Patho): p = 2.42e-102
A-D (Benign ~ Unif): p = 0.000124
A-D (Patho ~ Unif): p = 2.65e-73
Patient Data
Public Data
Uniform
cDNA position
B
General
Polarity
Acidity
Structure
Size
Origin
Amino Acid Position
C
>98% Conservation
Amino Acid Position
Property
Acidity
Origin
Size
General
Polarity
Structure
D
General
Polarity
Acidity
Structure
Size
Origin
Amino Acid Position

### SMARCC1

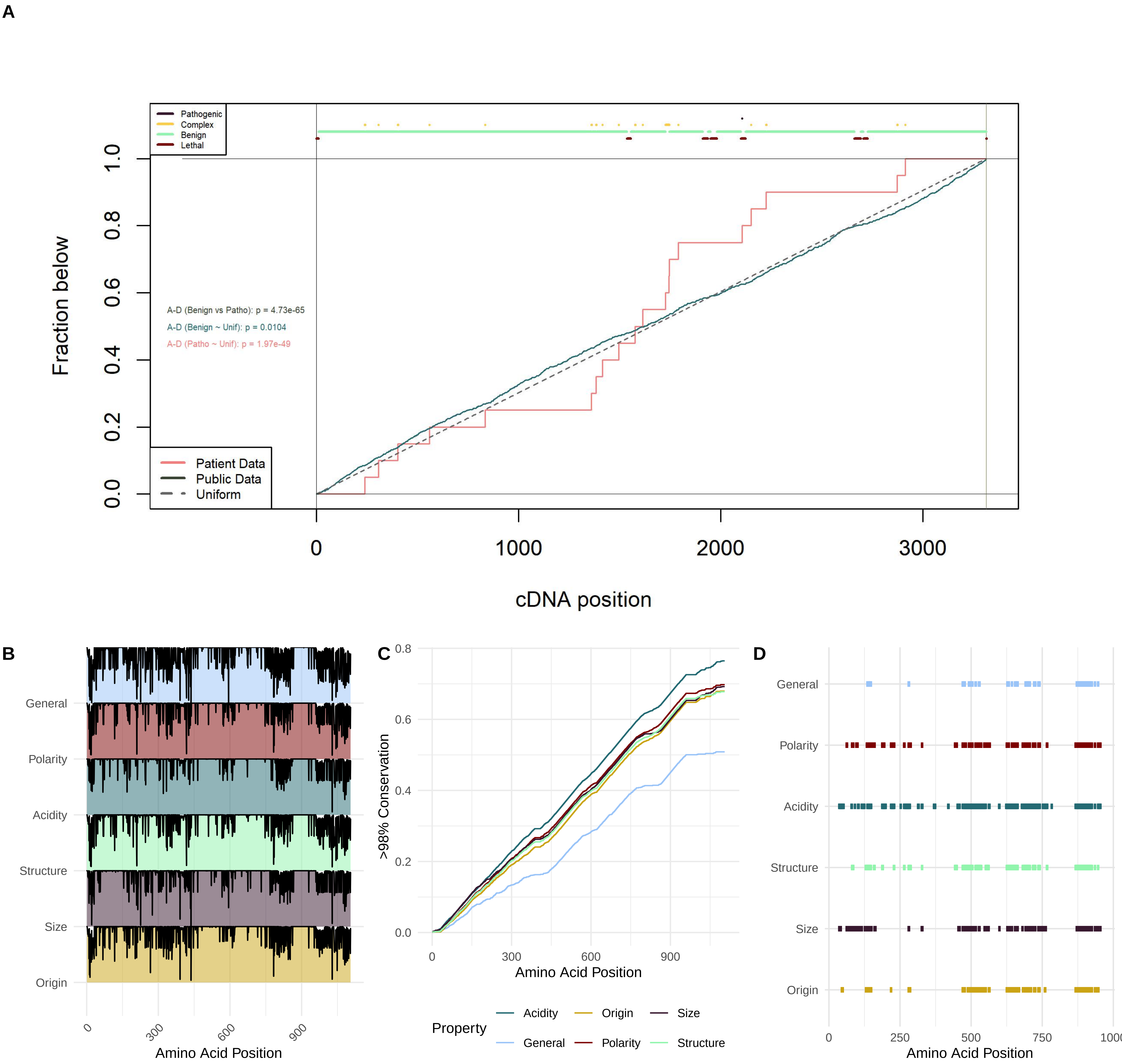
A
Pathogenic
Complex
Benign
Lethal
Fraction below
A-D (Benign vs Patho): p = 4.73e-65
A-D (Benign ~ Unif): p = 0.0104
A-D (Patho ~ Unif): p = 1.97e-49
Patient Data
Public Data
Uniform
cDNA position
B
General
Polarity
Acidity
Structure
Size
Origin
Amino Acid Position
C
>98% Conservation
Amino Acid Position
Property
Acidity
Origin
Size
General
Polarity
Structure
D
General
Polarity
Acidity
Structure
Size
Origin
Amino Acid Position

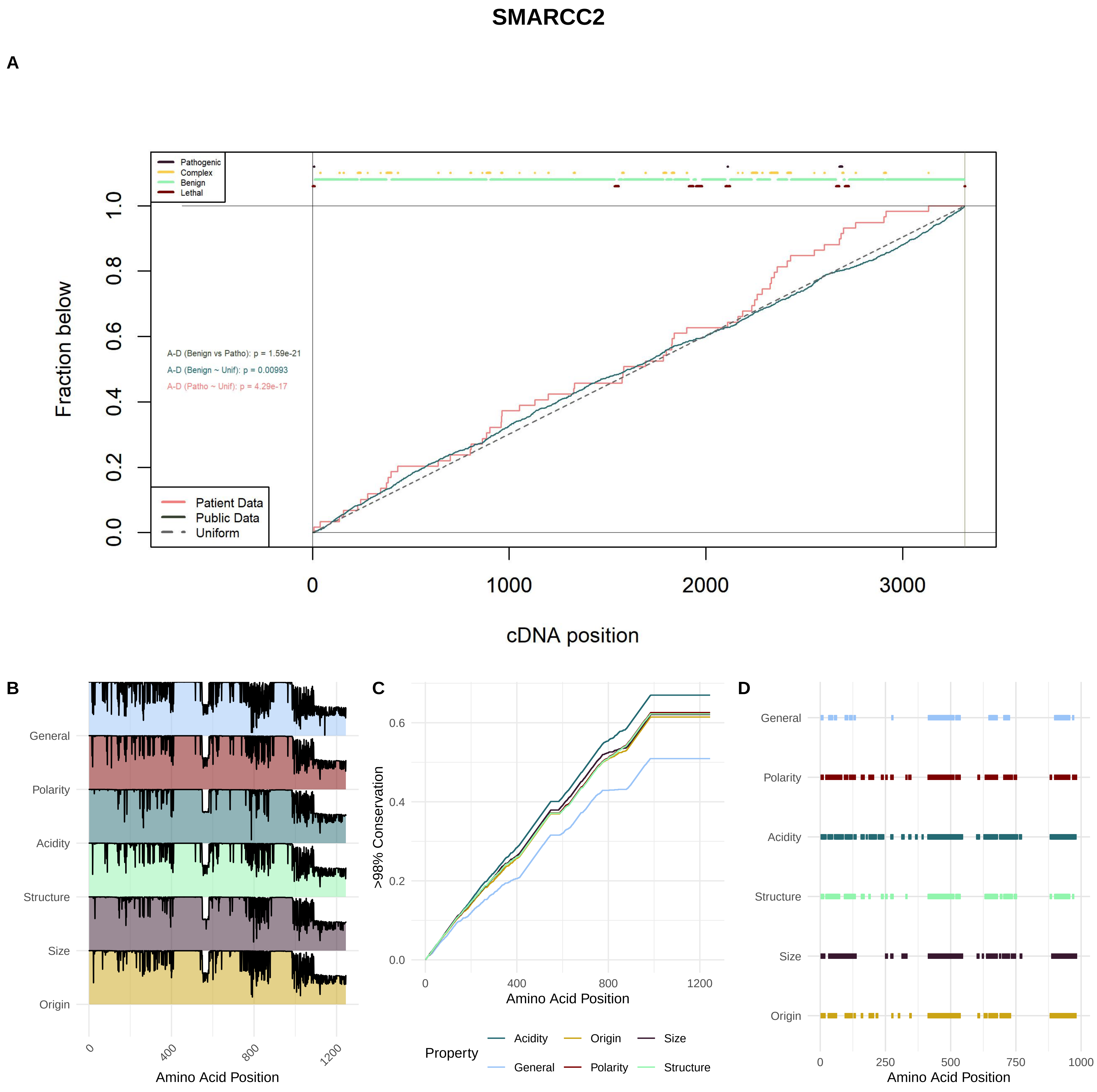

SMARCC2
A
Pathogenic
Complex
Benign
Lethal
Fraction below
A-D (Benign vs Patho): p = 1.59e-21
A-D (Benign ~ Unif): p = 0.00993
A-D (Patho ~ Unif): p = 4.29e-17
Patient Data
Public Data
Uniform
cDNA position
B
General
Polarity
Acidity
Structure
Size
Origin
Amino Acid Position
C
>98% Conservation
Amino Acid Position
Property
Acidity
Origin
Size
General
Polarity
Structure
D
General
Polarity
Acidity
Structure
Size
Origin
Amino Acid Position

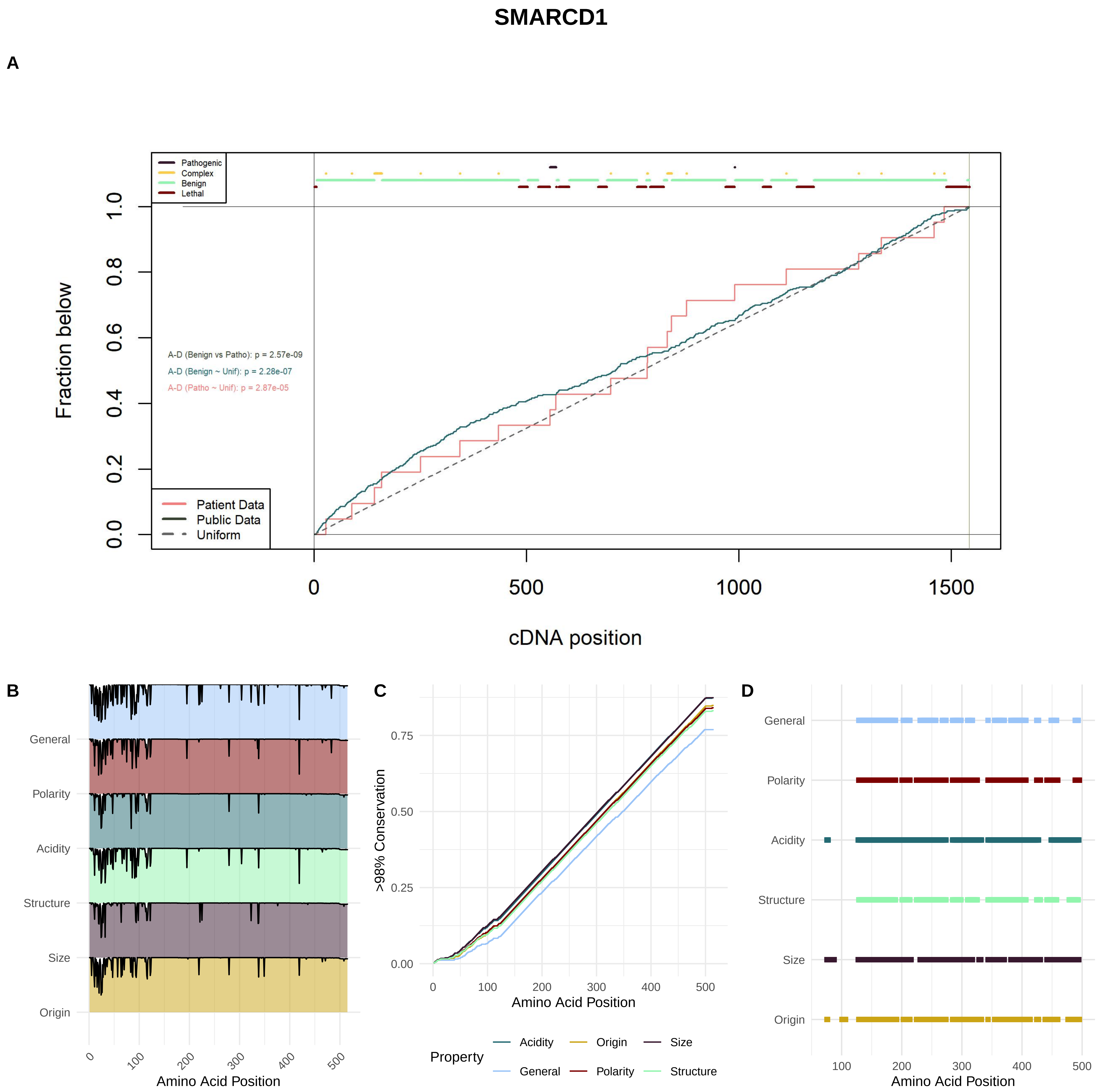
SMARCD1
A
Pathogenic
Complex
Benign
Lethal
Fraction below
A-D (Benign vs Patho): p = 2.57e-09
A-D (Benign ~ Unif): p = 2.28e-07
A-D (Patho ~ Unif): p = 2.87e-05
Patient Data
Public Data
Uniform
cDNA position
B
General
Polarity
Acidity
Structure
Size
Origin
Amino Acid Position
C
>98% Conservation
Amino Acid Position
Property
Acidity
Origin
Size
General
Polarity
Structure
D
General
Polarity
Acidity
Structure
Size
Origin
Amino Acid Position

### SMARCD2

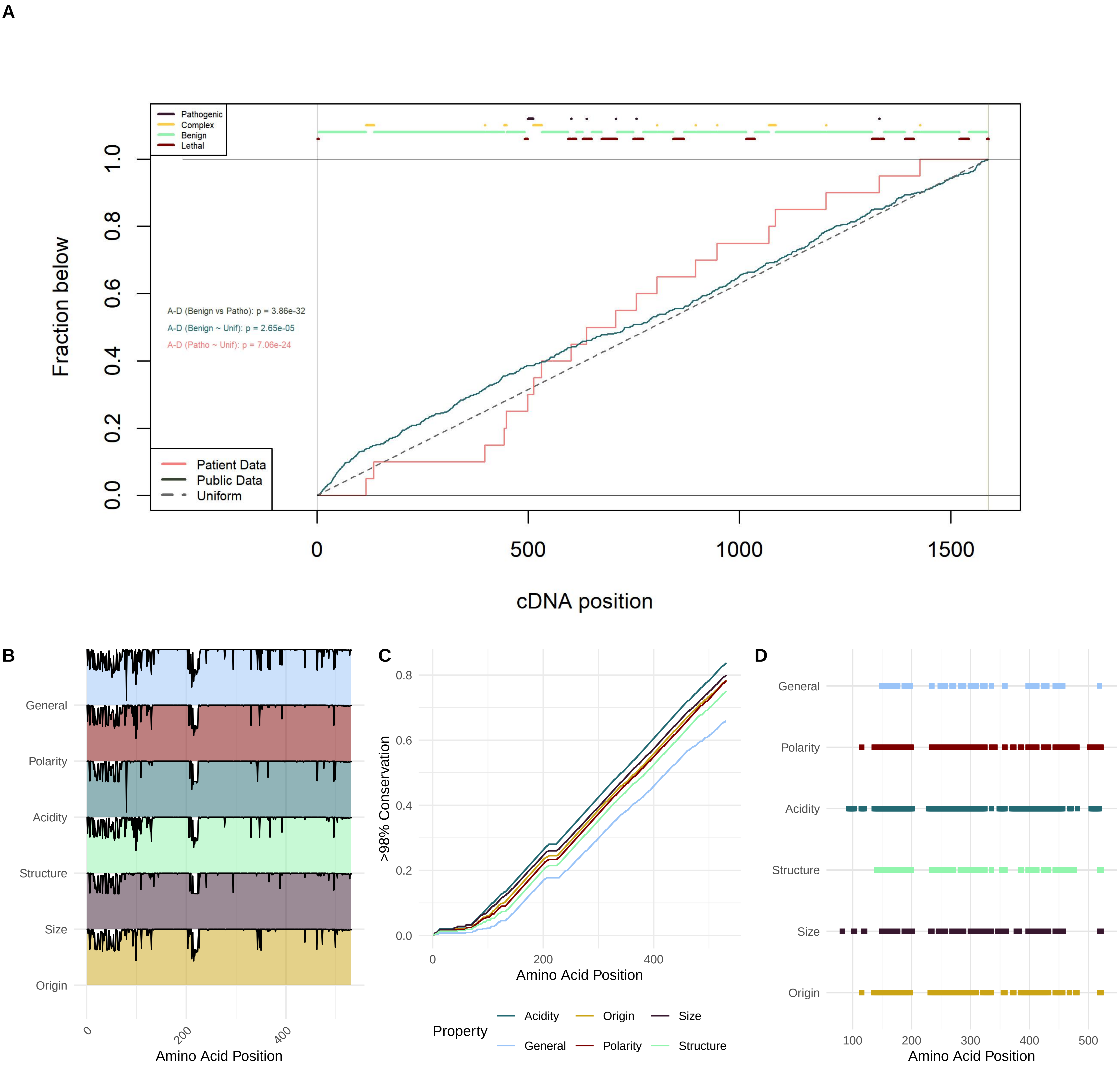

A
Pathogenic
Complex
Benign
Lethal
A-D (Benign vs Patho): p = 3.86e-32
A-D (Benign ~ Unif): p = 2.65e-05
A-D (Patho ~ Unif): p = 7.06e-24
Patient Data
Public Data
Uniform
Fraction below
cDNA position
B
General
Polarity
Acidity
Structure
Size
Origin
Amino Acid Position
C
>98% Conservation
Amino Acid Position
Property
Acidity
Origin
Size
General
Polarity
Structure
D
General
Polarity
Acidity
Structure
Size
Origin
Amino Acid Position

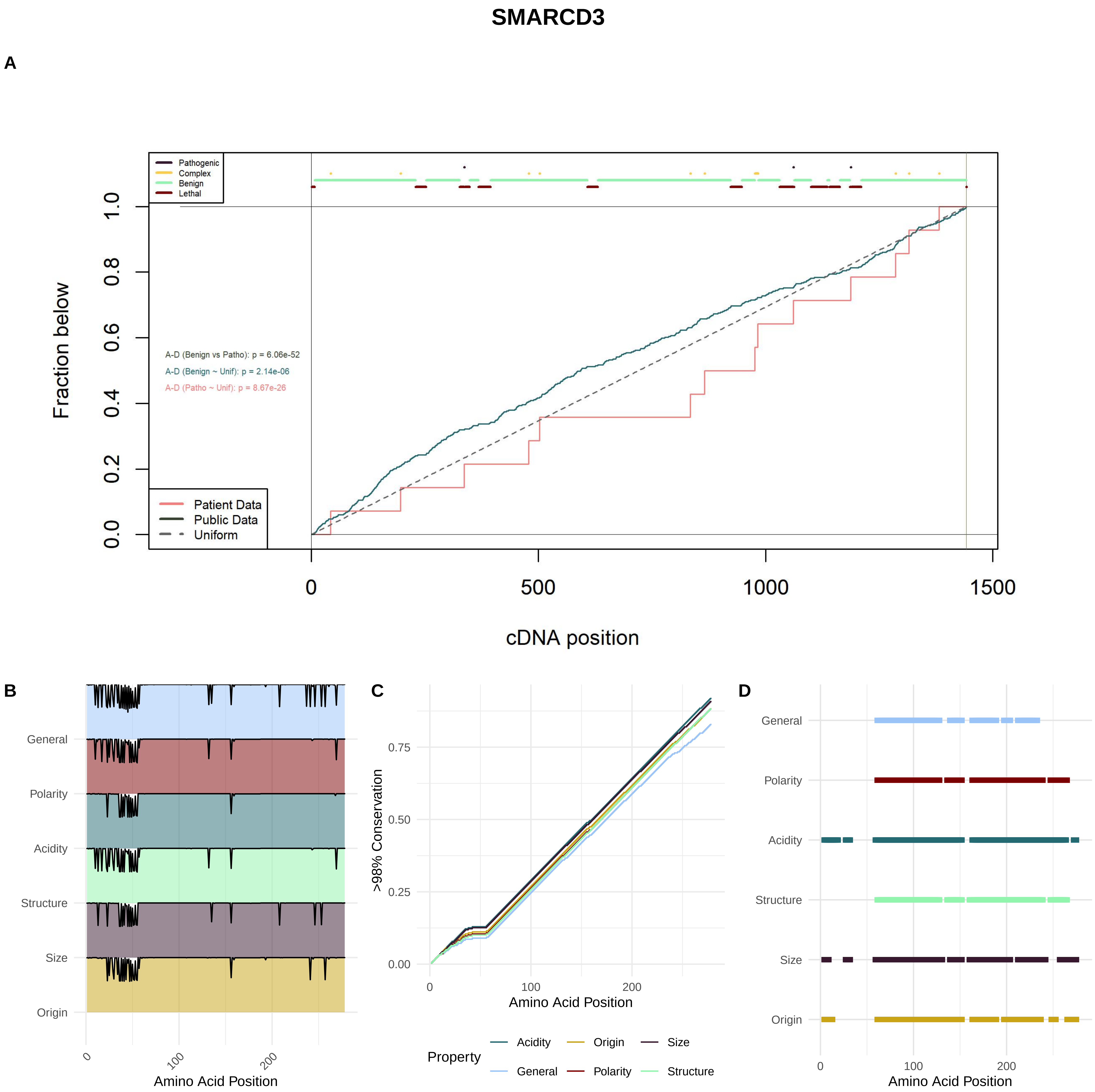
SMARCD3
A
Pathogenic
Complex
Benign
Lethal
Fraction below
0.0
0.2
0.4
0.6
0.8
1.0
A-D (Benign vs Patho): p = 6.06e-52
A-D (Benign ~ Unif): p = 2.14e-06
A-D (Patho ~ Unif): p = 8.67e-26
Patient Data
Public Data
Uniform
0
500
1000
1500
cDNA position
B
General
Polarity
Acidity
Structure
Size
Origin
0
100
200
Amino Acid Position
C
>98% Conservation
0.00
0.25
0.50
0.75
0
100
200
Amino Acid Position
Property
Acidity
Origin
Size
General
Polarity
Structure
D
General
Polarity
Acidity
Structure
Size
Origin
0
100
200
Amino Acid Position

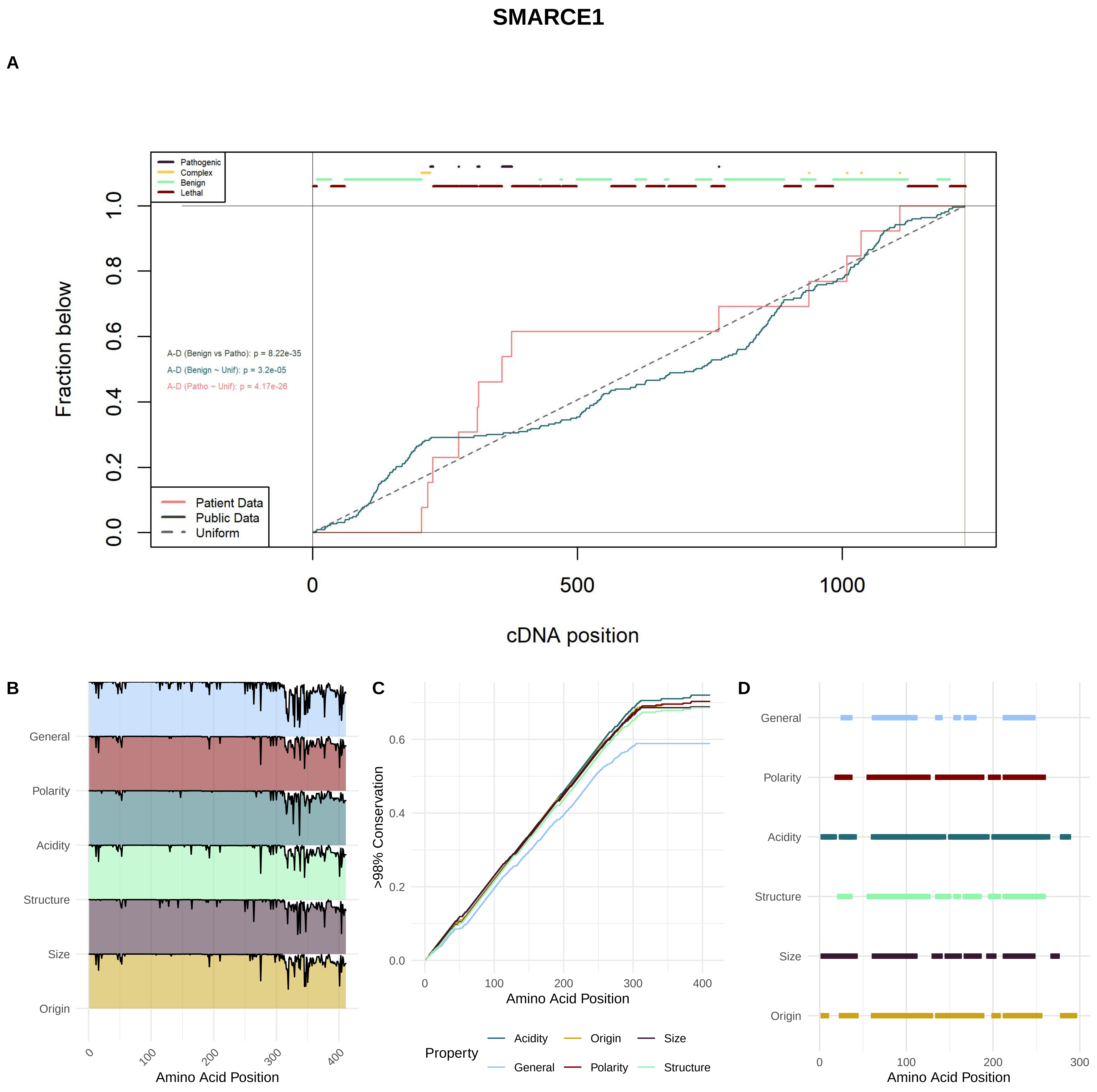

SMARCE1
A
Pathogenic
Complex
Benign
Lethal
Fraction below
A-D (Benign vs Patho): p = 8.22e-35
A-D (Benign ~ Unif): p = 3.2e-05
A-D (Patho ~ Unif): p = 4.17e-26
Patient Data
Public Data
Uniform
cDNA position
B
General
Polarity
Acidity
Structure
Size
Origin
Amino Acid Position
C
>98% Conservation
Amino Acid Position
Property
Acidity
Origin
Size
General
Polarity
Structure
D
General
Polarity
Acidity
Structure
Size
Origin
Amino Acid Position

### DPF1

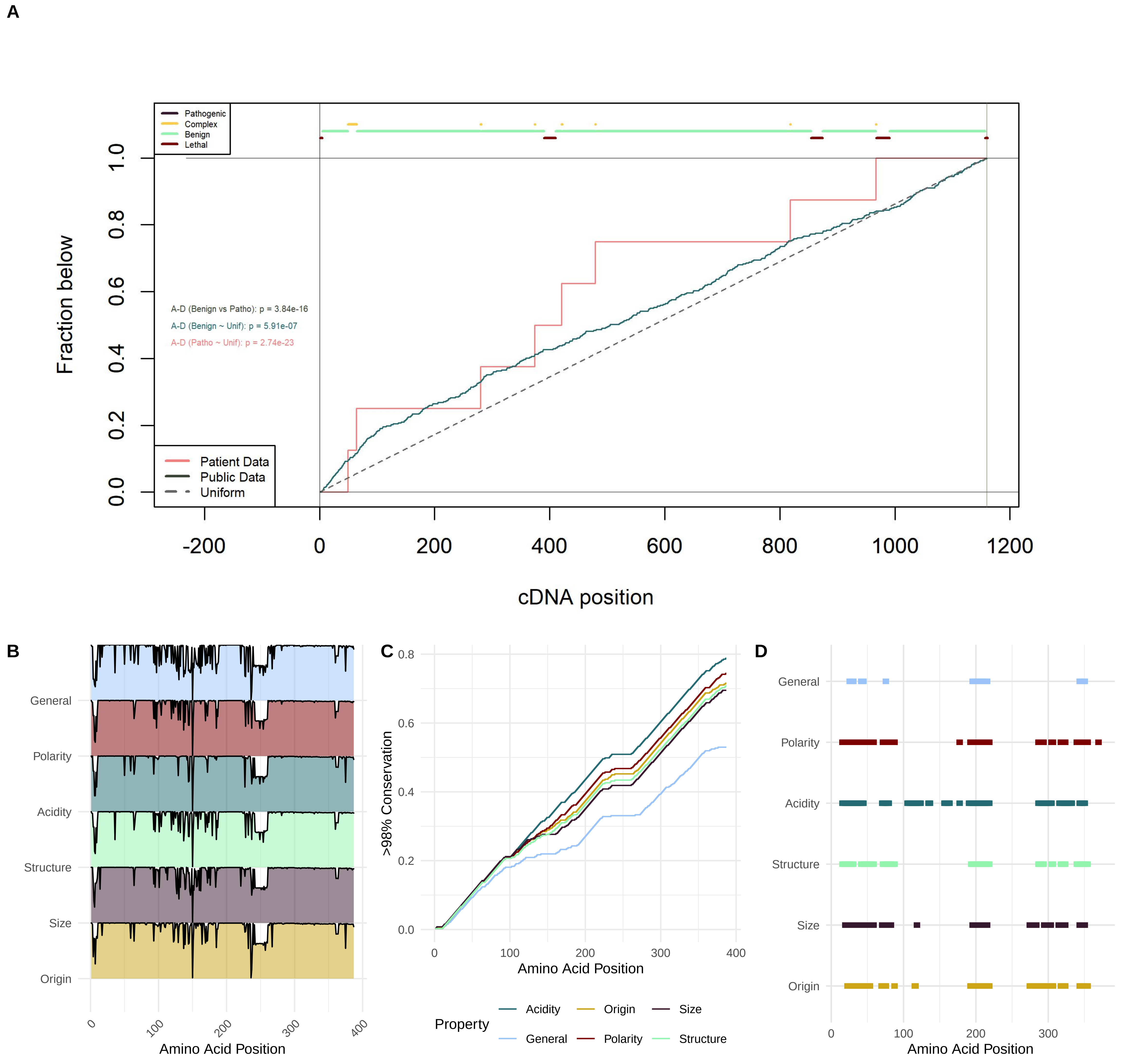
A
Pathogenic
Complex
Benign
Lethal
Fraction below
0.0
0.2
0.4
0.6
0.8
1.0
A-D (Benign vs Patho): p = 3.84e-16
A-D (Benign ~ Unif): p = 5.91e-07
A-D (Patho ~ Unif): p = 2.74e-23
Patient Data
Public Data
Uniform
-200
0
200
400
600
800
1000
1200
cDNA position
B
General
Polarity
Acidity
Structure
Size
Origin
0
100
200
300
400
Amino Acid Position
C
0.8
0.6
0.4
0.2
0.0
>98% Conservation
0
100
200
300
400
Amino Acid Position
Property
Acidity
Origin
Size
General
Polarity
Structure
D
General
Polarity
Acidity
Structure
Size
Origin
0
100
200
300
Amino Acid Position

### DPF2

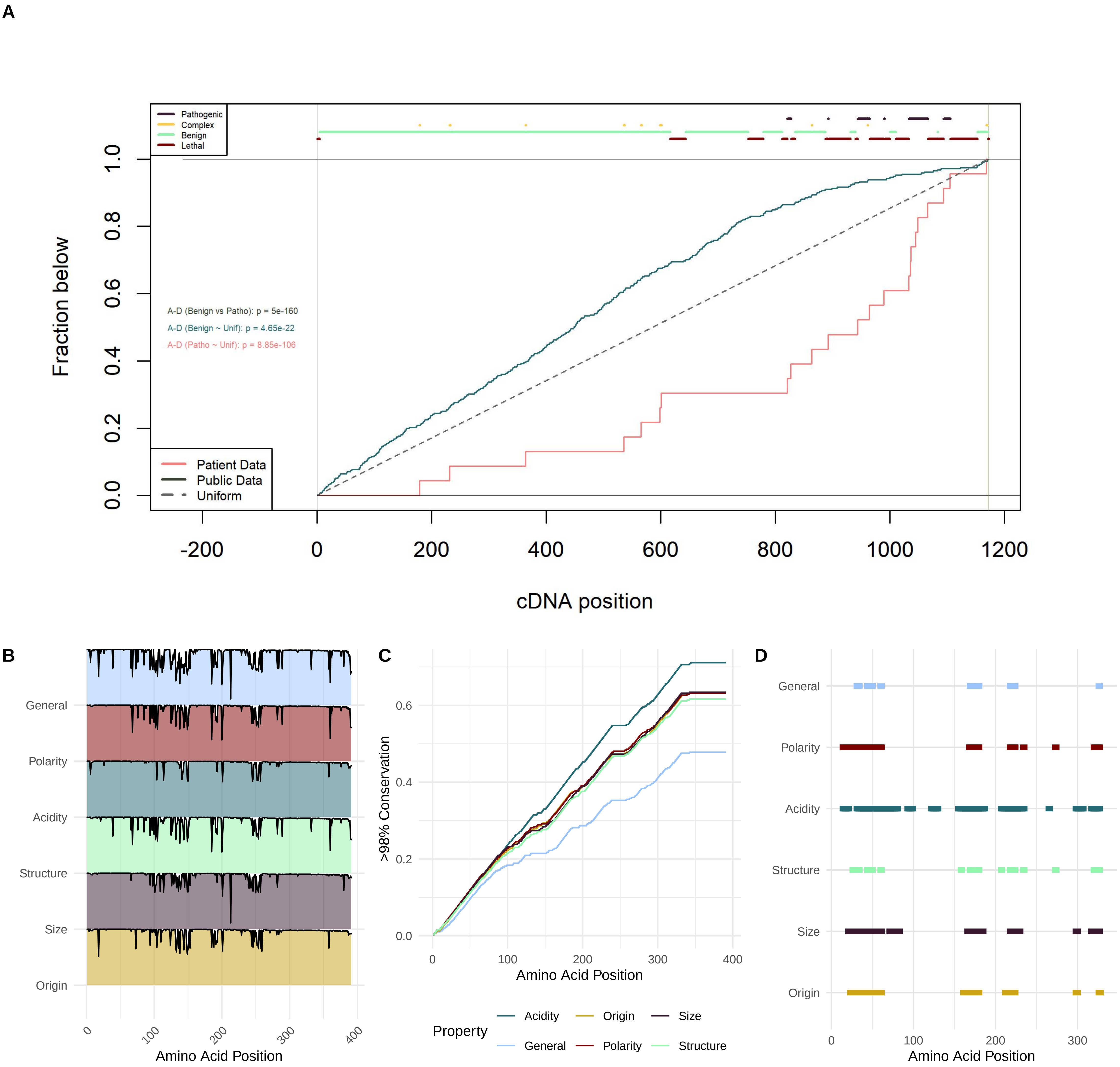

A
Pathogenic
Complex
Benign
Lethal
Fraction below
A-D (Benign vs Patho): p = 5e-160
A-D (Benign ~ Unif): p = 4.65e-22
A-D (Patho ~ Unif): p = 8.85e-106
Patient Data
Public Data
Uniform
cDNA position
B
General
Polarity
Acidity
Structure
Size
Origin
Amino Acid Position
C
>98% Conservation
Amino Acid Position
Property
Acidity
Origin
Size
General
Polarity
Structure
D
General
Polarity
Acidity
Structure
Size
Origin
Amino Acid Position

### DPF3

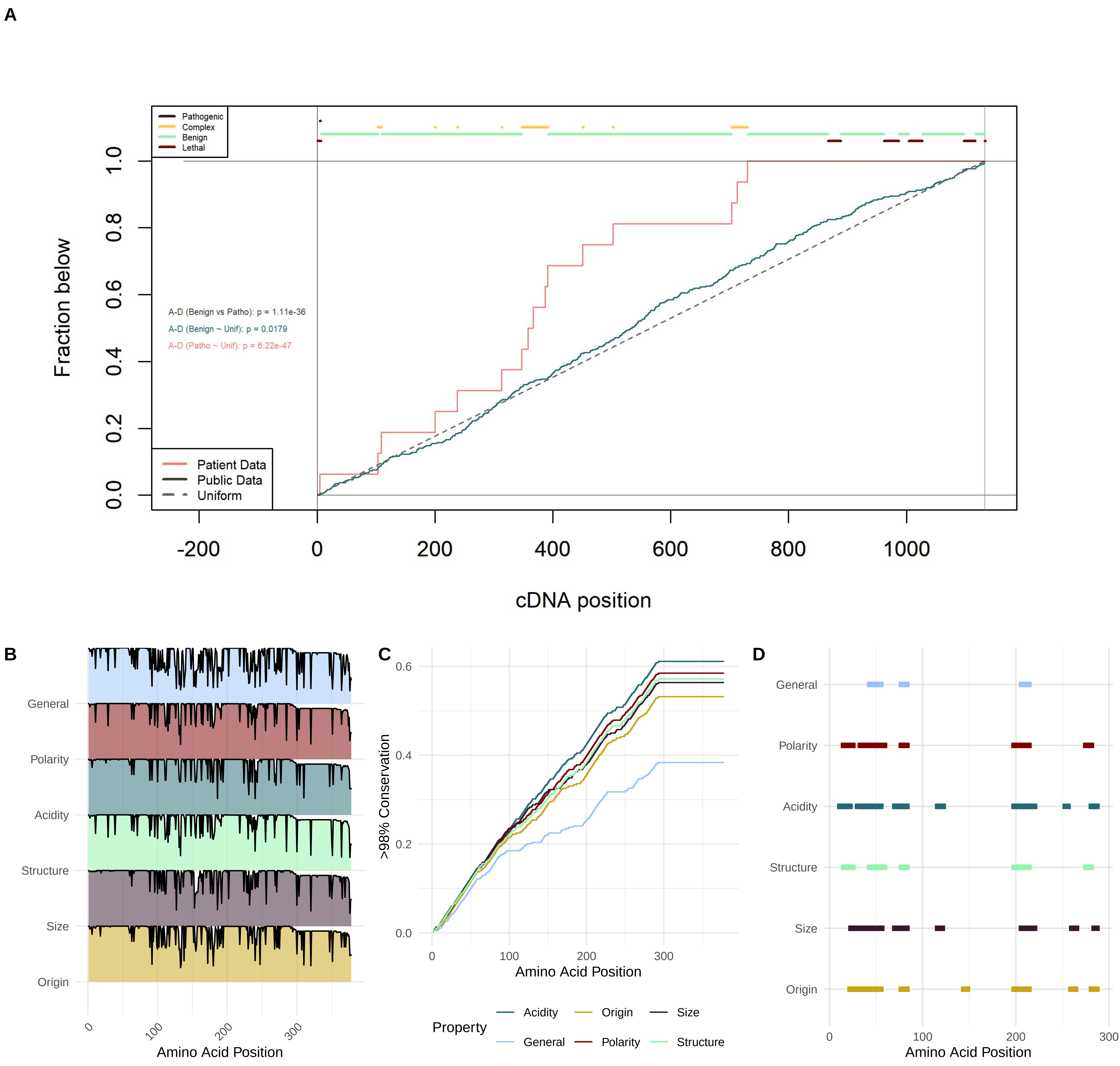
A
Pathogenic
Complex
Benign
Lethal
A-D (Benign vs Patho): p = 1.11e-36
A-D (Benign ~ Unif): p = 0.0179
A-D (Patho ~ Unif): p = 6.22e-47
Patient Data
Public Data
Uniform
Fraction below
0.0
0.2
0.4
0.6
0.8
1.0
-200
0
200
400
600
800
1000
cDNA position
B
General
Polarity
Acidity
Structure
Size
Origin
0
100
200
300
Amino Acid Position
C
0.0
0.2
0.4
0.6
>98% Conservation
0
100
200
300
Amino Acid Position
Property
Acidity
Origin
Size
General
Polarity
Structure
D
General
Polarity
Acidity
Structure
Size
Origin
0
100
200
300
Amino Acid Position

### ARID1A

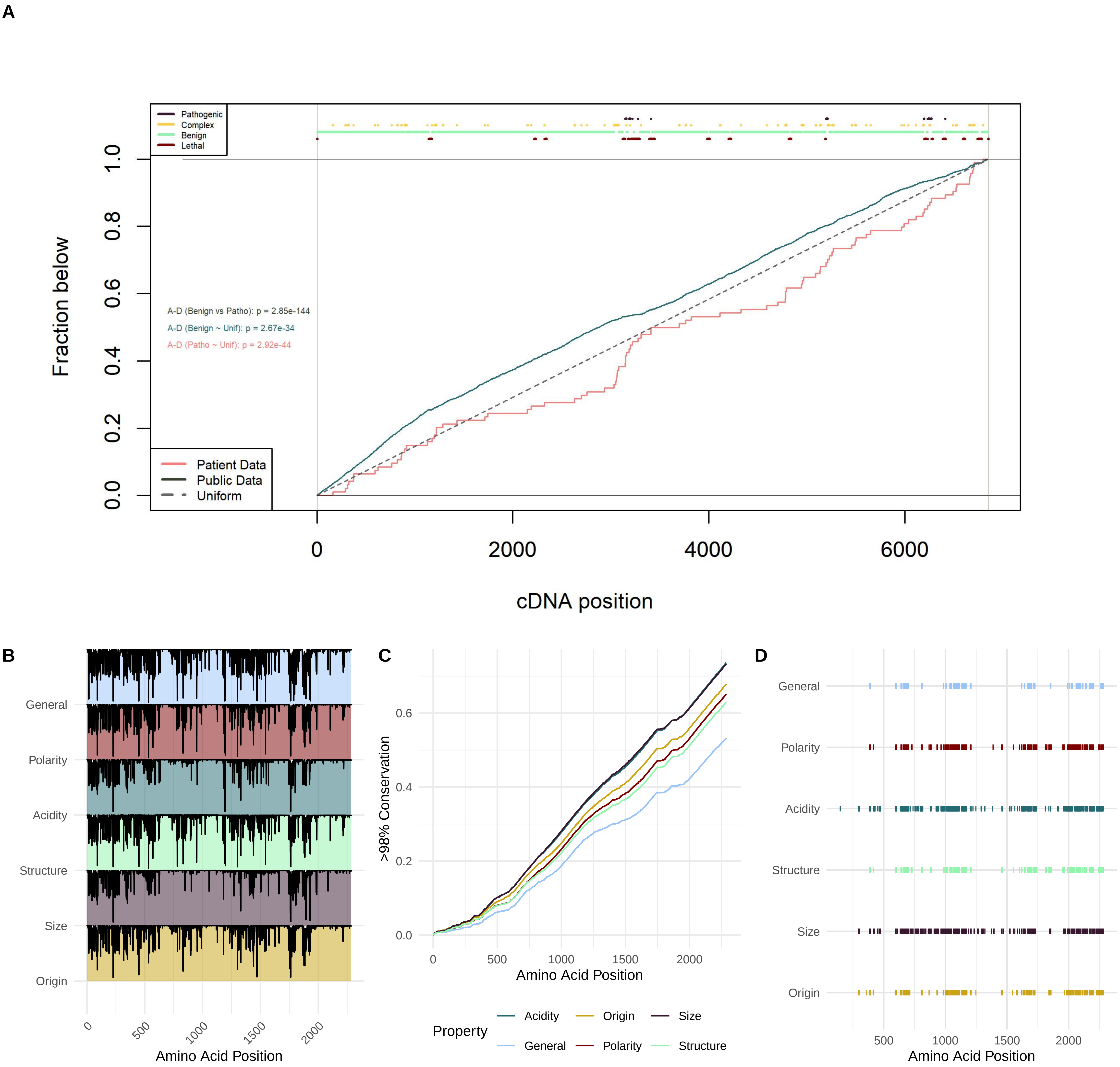

A
Pathogenic
Complex
Benign
Lethal
A-D (Benign vs Patho): p = 2.85e-144
A-D (Benign ~ Unif): p = 2.67e-34
A-D (Patho ~ Unif): p = 2.92e-44
Patient Data
Public Data
Uniform
Fraction below
0.0
0.2
0.4
0.6
0.8
1.0
0
2000
4000
6000
cDNA position
B
General
Polarity
Acidity
Structure
Size
Origin
0
500
1000
1500
2000
Amino Acid Position
C
>98% Conservation
0.0
0.2
0.4
0.6
0
500
1000
1500
2000
Amino Acid Position
Property
Acidity
Origin
Size
General
Polarity
Structure
D
General
Polarity
Acidity
Structure
Size
Origin
500
1000
1500
2000
Amino Acid Position

ARID1B

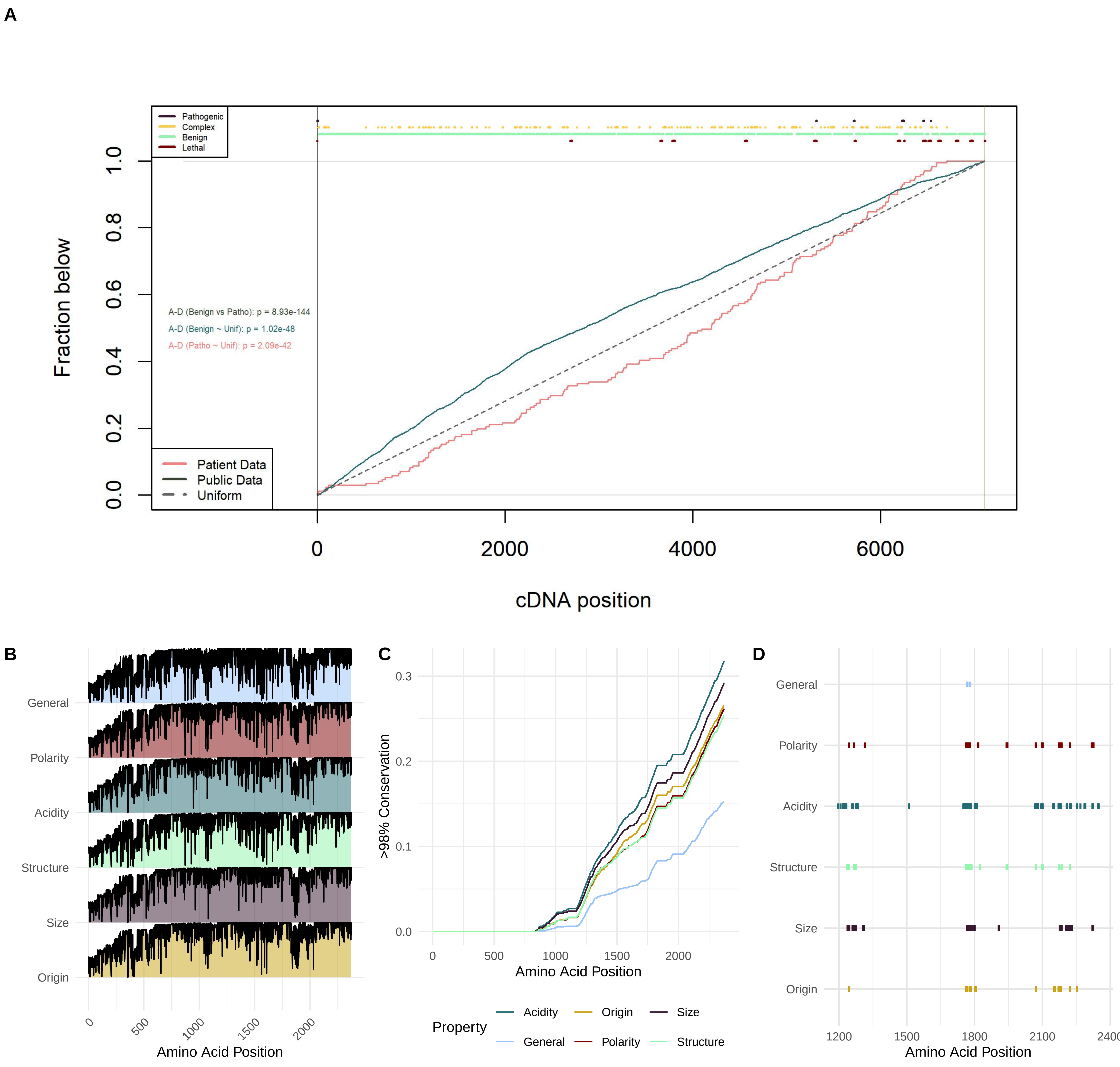

A
Pathogenic
Complex
Benign
Lethal
Fraction below
A-D (Benign vs Patho): p = 8.93e-144
A-D (Benign ~ Unif): p = 1.02e-48
A-D (Patho ~ Unif): p = 2.09e-42
Patient Data
Public Data
Uniform
cDNA position
B
General
Polarity
Acidity
Structure
Size
Origin
Amino Acid Position
C
>98% Conservation
Amino Acid Position
Property
Acidity
Origin
Size
General
Polarity
Structure
D
General
Polarity
Acidity
Structure
Size
Origin
Amino Acid Position

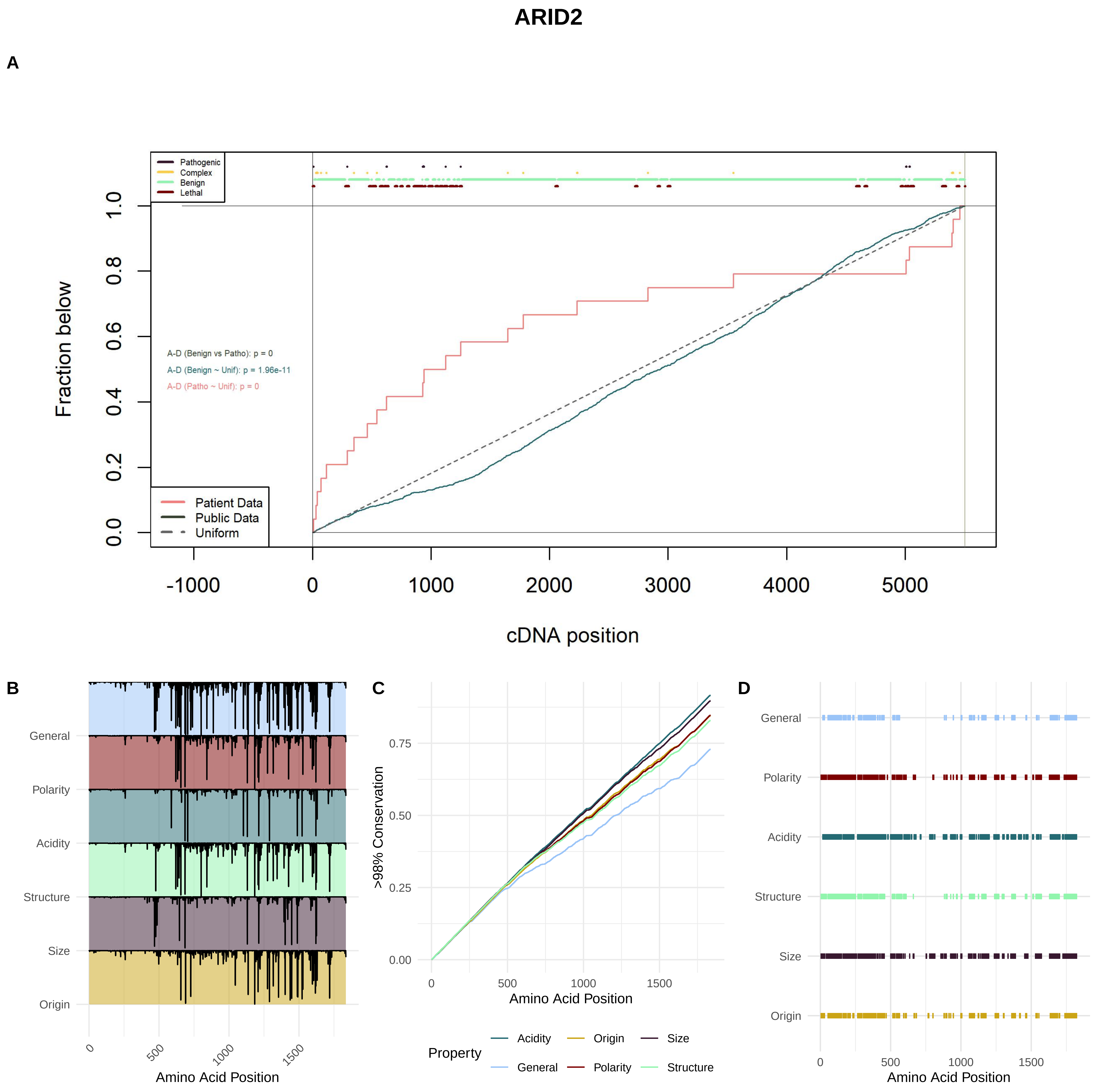

ARID2
A
Pathogenic
Complex
Benign
Lethal
Fraction below
0.0
0.2
0.4
0.6
0.8
1.0
A-D (Benign vs Patho): p = 0
A-D (Benign ~ Unif): p = 1.96e-11
A-D (Patho ~ Unif): p = 0
Patient Data
Public Data
Uniform
-1000
0
1000
2000
3000
4000
5000
cDNA position
B
General
Polarity
Acidity
Structure
Size
Origin
0
500
1000
1500
Amino Acid Position
C
>98% Conservation
0.00
0.25
0.50
0.75
0
500
1000
1500
Amino Acid Position
Property
Acidity
Origin
Size
General
Polarity
Structure
D
General
Polarity
Acidity
Structure
Size
Origin
0
500
1000
1500
Amino Acid Position
