## Supplementary Tables for "Gene Specific Pathogenicity Predictor for Chromatin-Remodeling BAF Complex-Associated Neurodevelopmental Disorders"

**Table S1. BAF Subunits Included in Generalization**

| Subunit | Gene |
|---|---|
| SMARC | *SMARCB1* |
| | *SMARCC1* |
| | *SMARCC2* |
| | *SMARCD1* |
| | *SMARCD2* |
| | *SMARCD3* |
| | *SMARCE1* |
| DPF | *DPF1* |
| | *DPF2* |
| | *DPF3* |
| ARID | *ARID1A* |
| | *ARID1B* |
| | *ARID2* |

**Table S2. Performance metrics of alternative pathogenicity predictors.**

| | SIFT | EVE | AM | CLINPRED | REVEL>0.75 | REVEL>0.5 | PolyPhen | | |
|---|---|---|---|---|---|---|---|---|---|
| Accuracy | 0.254 | 0.826 | 0.620 | 0.341 | 0.859 | 0.698 | 0.510 | | |
| Sensitivity | 0.840 | 0.507 | 0.743 | 0.836 | 0.477 | 0.721 | 0.819 | | |
| Specificity | 0.950 | 0.920 | 0.926 | 0.908 | 0.916 | 0.924 | 0.893 | | |
| AUROC Max | 0.846 | 0.619 | 0.856 | 0.720 | 0.920 | 0.863 | 0.704 | | |
| AUROC Min | 0.556 | 0.512 | 0.592 | 0.536 | 0.720 | 0.601 | 0.537 | | |
| AUROC Est | 0.701 | 0.566 | 0.724 | 0.628 | 0.820 | 0.732 | 0.621 | | |
| **Generalization** | | | | | | | | | |
| | SIFT | EVE | AM | CLINPRED | REVEL>0.75 | REVEL>0.5 | PolyPhen | CADD | BAF_Wald |
| Accuracy | 0.469 | 0.707 | 0.607 | 0.509 | 0.613 | 0.587 | 0.545 | 0.654 | 0.630 |
| Sensitivity | 0.831 | 0.556 | 0.633 | 0.754 | 0.073 | 0.263 | 0.471 | 0.153 | 0.173 |
| Specificity | 0.290 | 0.826 | 0.592 | 0.398 | 0.943 | 0.787 | 0.579 | 0.906 | 0.999 |
| AUROC Max | 0.709 | 0.845 | 0.701 | 0.691 | 0.613 | 0.596 | 0.553 | 0.693 | NA |
| AUROC Min | 0.561 | 0.691 | 0.613 | 0.576 | 0.508 | 0.525 | 0.525 | 0.530 | NA |
| AUROC Est | 0.635 | 0.768 | 0.657 | 0.633 | 0.561 | 0.560 | 0.539 | 0.611 | 0.762 |

**Table S3. Preprocessing steps included for each workflow.**

| Workflow | Dummy Variable | Remove Single Instance | Impute | Decorrelate | Normalize | Transform |
|---|---|---|---|---|---|---|
| No SMOTE Bagged Tree | Yes | Yes | Yes | Yes | Yes | No |
| No SMOTE Balanced Random Forest | Yes | Yes | Yes | Yes | Yes | No |
| No SMOTE Bayes | Yes | Yes | Yes | Yes | Yes | No |
| No SMOTE Random Forest | Yes | Yes | Yes | Yes | Yes | No |
| No SMOTE Boosted Tree | Yes | Yes | Yes | Yes | Yes | No |
| No SMOTE Weighted Logistic Regression | Yes | Yes | Yes | Yes | Yes | No |
| SMOTE kNN | Yes | Yes | Yes | Yes | Yes | Yes |
| SMOTE Logistic Regression | Yes | Yes | Yes | Yes | Yes | Yes |
| SMOTE Neural Network | Yes | Yes | Yes | Yes | Yes | Yes |
| SMOTE Random Forest | Yes | Yes | Yes | Yes | Yes | Yes |
| SMOTE Tree | Yes | Yes | Yes | Yes | Yes | Yes |
| Support Vector Machine Linear | Yes | Yes | Twice | Yes | Yes | No |
| Support Vector Machine Polynomial | Yes | Yes | Twice | Yes | Yes | No |
| Support Vector Machine Radial Basis Function | Yes | Yes | Twice | Yes | Yes | No |
| Support Vector Machine Sigmoid | Yes | Yes | Twice | Yes | Yes | No |

**Table S4. Performance metrics of each workflow with *SMARCA2* and *SMARCA4* .**

| Workflow | Mean Accuracy | Mean AUROC | Mean Sensitivity | Mean Specificity | Standard Deviation Accuracy | Standard Deviation AUROC | Standard Deviation Sensitivity | Standard Deviation Specificity |
|---|---|---|---|---|---|---|---|---|
| No SMOTE Bagged Tree | 0.914 | 0.931 | 0.958 | 0.663 | 0.015 | 0.028 | 0.011 | 0.063 |
| No SMOTE Balanced Random Forest | 0.909 | 0.940 | 0.966 | 0.585 | 0.019 | 0.041 | 0.019 | 0.093 |
| No SMOTE Bayes | 0.880 | 0.872 | 0.924 | 0.634 | 0.016 | 0.020 | 0.019 | 0.052 |
| No SMOTE Random Forest | 0.909 | 0.940 | 0.966 | 0.585 | 0.019 | 0.041 | 0.019 | 0.093 |
| No SMOTE Boosted Tree | 0.907 | 0.951 | 0.955 | 0.634 | 0.015 | 0.012 | 0.008 | 0.063 |
| No SMOTE Weighted Logistic Regression | 0.908 | 0.953 | 0.958 | 0.622 | 0.017 | 0.009 | 0.013 | 0.024 |
| SMOTE kNN | 0.869 | 0.898 | 0.884 | 0.785 | 0.019 | 0.011 | 0.024 | 0.044 |
| SMOTE Logistic Regression | 0.887 | 0.951 | 0.890 | 0.867 | 0.010 | 0.008 | 0.022 | 0.075 |
| SMOTE Neural Network | 0.886 | 0.917 | 0.911 | 0.746 | 0.015 | 0.029 | 0.025 | 0.083 |
| SMOTE Random Forest | 0.904 | 0.952 | 0.937 | 0.714 | 0.014 | 0.009 | 0.012 | 0.045 |
| SMOTE Tree | 0.869 | 0.916 | 0.882 | 0.797 | 0.026 | 0.016 | 0.028 | 0.030 |
| Support Vector Machine Linear | 0.868 | 0.949 | 0.862 | 0.905 | 0.031 | 0.008 | 0.046 | 0.069 |
| Support Vector Machine Polynomial | 0.692 | 0.820 | 0.660 | 0.874 | 0.160 | 0.222 | 0.197 | 0.106 |
| Support Vector Machine Radial Basis Function | 0.721 | 0.799 | 0.705 | 0.812 | 0.132 | 0.218 | 0.172 | 0.174 |
| Support Vector Machine Sigmoid | 0.709 | 0.792 | 0.678 | 0.887 | 0.109 | 0.239 | 0.126 | 0.077 |

**Table S5. Results from tuning Random Forest.**

| Number of Features | Trees | Minimun Nodes | Configuration |
|---|---|---|---|
| 4 | 1143 | 3 | Preprocessor 1 Model 05 |

**Table S6. Performance on different datasets.**

| Metric | Estimator | Estimate | Dataset |
|---|---|---|---|
| Accuracy | Binary | 1.000 | Training |
| Accuracy | Binary | 0.937 | Test |
| Accuracy | Binary | 0.933 | Holdout |

**Table S7. Performance of predictor on other BAF subunits.**

| Metric | Estimator | Estimate | Dataset |
|---|---|---|---|
| Accuracy | Binary | 0.645 | ARID |
| Cohen's Kappa | Binary | 0.004 | ARID |
| AUROC | Binary | 0.552 | ARID |
| Accuracy | Binary | 0.843 | DPF |
| Cohen's Kappa | Binary | 0.121 | DPF |
| AUROC | Binary | 0.598 | DPF |
| Accuracy | Binary | 0.577 | SMARC |
| Cohen's Kappa | Binary | 0.067 | SMARC |
| AUROC | Binary | 0.541 | SMARC |
| Accuracy | Binary | 0.685 | Combined |
| Cohen's Kappa | Binary | 0.066 | Combined |
| AUROC | Binary | 0.559 | Combined |

**Table S8. Workflow performance after retraining with all BAF subunits.**

| Workflow | Mean Accuracy | Mean AUROC | Mean Sensitivity | Mean Specificity | Standard Deviation Accuracy | Standard Deviation AUROC | Standard Deviation Sensitivity | Standard Deviation Specificity |
|---|---|---|---|---|---|---|---|---|
| No SMOTE Bagged Tree | 0.880 | 0.887 | 0.937 | 0.665 | 0.012 | 0.028 | 0.017 | 0.042 |
| No SMOTE Balanced Random Forest | 0.888 | 0.904 | 0.965 | 0.598 | 0.010 | 0.024 | 0.011 | 0.057 |
| No SMOTE Bayes | 0.796 | 0.778 | 0.861 | 0.562 | 0.055 | 0.010 | 0.081 | 0.046 |
| No SMOTE Random Forest | 0.888 | 0.904 | 0.965 | 0.599 | 0.010 | 0.024 | 0.011 | 0.056 |
| No SMOTE Boosted Tree | 0.876 | 0.905 | 0.951 | 0.595 | 0.011 | 0.022 | 0.012 | 0.064 |
| No SMOTE Weighted Logistic Regression | 0.861 | 0.864 | 0.960 | 0.493 | 0.013 | 0.032 | 0.018 | 0.034 |
| SMOTE kNN | 0.799 | 0.818 | 0.820 | 0.717 | 0.014 | 0.015 | 0.021 | 0.030 |
| SMOTE Logistic Regression | 0.796 | 0.862 | 0.815 | 0.722 | 0.037 | 0.033 | 0.046 | 0.079 |
| SMOTE Neural Network | 0.795 | 0.847 | 0.814 | 0.722 | 0.033 | 0.032 | 0.045 | 0.061 |
| SMOTE Random Forest | 0.868 | 0.907 | 0.918 | 0.681 | 0.021 | 0.030 | 0.025 | 0.047 |
| SMOTE Tree | 0.823 | 0.851 | 0.854 | 0.703 | 0.023 | 0.026 | 0.041 | 0.068 |
| Support Vector Machine Linear | 0.797 | 0.858 | 0.815 | 0.726 | 0.031 | 0.030 | 0.043 | 0.065 |
| Support Vector Machine Polynomial | 0.674 | 0.613 | 0.651 | 0.760 | 0.092 | 0.284 | 0.129 | 0.059 |
| Support Vector Machine Radial Basis Function | 0.706 | 0.604 | 0.707 | 0.704 | 0.101 | 0.291 | 0.154 | 0.118 |
| Support Vector Machine Sigmoid | 0.672 | 0.560 | 0.650 | 0.756 | 0.087 | 0.298 | 0.123 | 0.059 |

**Table S9. Performance on training, test, and holdout sets with all BAF subunits.**

| Metric | Estimator | Estimate | Dataset |
|---|---|---|---|
| Accuracy | Binary | 0.998 | Training |
| Cohen's Kappa | Binary | 0.994 | Training |
| Accuracy | Binary | 0.9032 | Test |
| Cohen's Kappa | Binary | 0.6343 | Test |
| Accuracy | Binary | 0.8988 | Holdout |
| Cohen's Kappa | Binary | 0.6757 | Holdout |

**Table S10. Performance metrics when "pathogenic window" feature is removed.**

| Workflow | Mean Accuracy | Mean AUROC | Mean Sensitivity | Mean Specificity | Standard Deviation Accuracy | Standard Deviation AUROC | Standard Deviation Sensitivity | Standard Deviation Specificity |
|---|---|---|---|---|---|---|---|---|
| No SMOTE Bagged Tree | 0.906 | 0.950 | 0.938 | 0.792 | 0.008 | 0.005 | 0.018 | 0.036 |
| No SMOTE Balanced Random Forest | 0.897 | 0.945 | 0.952 | 0.699 | 0.020 | 0.041 | 0.023 | 0.110 |
| No SMOTE Bayes | 0.835 | 0.876 | 0.877 | 0.683 | 0.020 | 0.012 | 0.037 | 0.075 |
| No SMOTE Random Forest | 0.897 | 0.945 | 0.952 | 0.699 | 0.020 | 0.041 | 0.023 | 0.110 |
| No SMOTE Boosted Tree | 0.904 | 0.961 | 0.943 | 0.764 | 0.006 | 0.006 | 0.018 | 0.040 |
| No SMOTE Weighted Logistic Regression | 0.872 | 0.916 | 0.940 | 0.627 | 0.008 | 0.011 | 0.011 | 0.040 |
| SMOTE kNN | 0.812 | 0.882 | 0.818 | 0.795 | 0.014 | 0.018 | 0.022 | 0.046 |
| SMOTE Logistic Regression | 0.810 | 0.913 | 0.810 | 0.811 | 0.008 | 0.011 | 0.018 | 0.058 |
| SMOTE Neural Network | 0.841 | 0.914 | 0.853 | 0.800 | 0.023 | 0.015 | 0.035 | 0.051 |
| SMOTE Random Forest | 0.905 | 0.959 | 0.929 | 0.821 | 0.008 | 0.005 | 0.019 | 0.040 |
| SMOTE Tree | 0.841 | 0.896 | 0.821 | 0.911 | 0.025 | 0.025 | 0.030 | 0.018 |
| Support Vector Machine Linear | 0.792 | 0.908 | 0.780 | 0.838 | 0.028 | 0.009 | 0.049 | 0.063 |
| Support Vector Machine Polynomial | 0.715 | 0.521 | 0.681 | 0.840 | 0.083 | 0.368 | 0.116 | 0.067 |
| Support Vector Machine Radial Basis Function | 0.715 | 0.585 | 0.710 | 0.738 | 0.126 | 0.353 | 0.200 | 0.237 |
| Support Vector Machine Sigmoid | 0.712 | 0.520 | 0.678 | 0.839 | 0.048 | 0.367 | 0.064 | 0.063 |

**Table S11. Performance on training, test, and holdout data when "pathogenic window" feature is removed.**

| Metric | Estimator | Estimate | Dataset |
|---|---|---|---|
| Accuracy | Binary | 0.998 | Training |
| Cohen's Kappa | Binary | 0.994 | Training |
| Accuracy | Binary | 0.933 | Test |
| Cohen's Kappa | Binary | 0.762 | Test |
| Accuracy | Binary | 0.929 | Holdout |
| Cohen's Kappa | Binary | 0.792 | Holdout |
